# STECode: an automated virulence barcode generator to aid clinical and public health risk assessment of Shiga toxin-producing *Escherichia coli*

**DOI:** 10.1101/2024.05.07.593058

**Authors:** Eby M. Sim, Winkie Fong, Carl Suster, Jessica E. Agius, Shona Chandra, Basel Suliman, Qinning Wang, Christine Ngo, Connor Finemore, Sharon C-A. Chen, Kerri Basile, Vitali Sintchenko

## Abstract

Shiga Toxin (Stx) producing *Escherichia coli* (STEC) is a subset of pathogenic *E. coli* that can produce two types of Stx, Stx1 and Stx2, which can be further subtyped into four and 15 subtypes respectively. Not all subtypes, however, are equal in virulence potential, and the risk of severe disease including haemolytic uraemic syndrome has been linked to certain Stx2 subtypes e.g. Stx2a, Stx2d, highlighting the importance to survey *stx* subtypes. Previously, we developed a STEC virulence barcode to capture pertinent information on virulence genes to infer pathogenic potential. However, the process required multiple manual curation steps to determine the barcode. Here we introduce STECode, a bioinformatic tool to automate the STEC virulence barcode generation from sequencing reads or genomic assemblies. The development, and validation of STECode is described using a set of publicly available completed STEC genomes, along with their corresponding short reads. STECode was applied to interrogate the virulence landscape and molecular epidemiology of human STEC isolated during the period of the international border closures related to COVID-19 in the state of New South Wales, Australia.

**Impact statement:** Whole genome sequencing has been used to great effect in the genomic surveillance of STEC for public health purposes via the tracking of outbreaks. With STECode, we present a method to generate a STEC virulence barcode which captures pertinent subtyping information, useful for genomic inference of pathogenic potential. A key blind spot generated in short-read sequencing is the inability to detect the presence of multiple, isogenic *stx* copies in STEC. STECode mitigates this by inferring and reporting on the possibility of this occurrence. We envisage that this tool will value-add current genomic surveillance workflows through the ability to infer pathogenic potential.

## Introduction

Shiga-toxin producing *Escherichia coli* (STEC) is a catch-all term to describe any *E. coli* that can produce Shiga toxin (Stx). Infections with STEC can cause different clinical syndromes ranging from asymptomatic carriage, acute gastroenteritis, progression to Haemolytic Ureamic Syndrome (HUS), neurological complications and death (1). Progression to the complication of HUS is considered to adversely impact clinical outcomes following an STEC infection (1). First described in *Shigella dysenteriae* (2), Stx is an A_1_B_5_ holotoxin consisting of an enzymatic active A subunit and a pentameric receptor-binding B subunit (3). The ability to produce Stx is conferred onto *E. coli* via by a prophage (*stx* prophage) (4). The genes that encode the A and B subunits form an operon, and are located downstream of the phage anti-terminator gene, linking toxin production to the induction of the *stx* prophage from its host genome (4). STEC produces two types of Stx, Stx1 and Stx2, with four Stx1 subtypes (Stx1a, Stx1c, Stx1d and Stx1e) and 15 Stx2 subtypes (Stx2a through to Stx2o) currently described (5–11). Progression to HUS following STEC infection is linked to the production of Stx, with the presence of Stx2a and Stx2d being strongly associated with severe outcomes (6, 7, 12).

Apart from Stx production, some STEC also harbour additional virulence factors which aid adhesion to the intestinal epithelial cells, and can aggravate the clinical outcomes. Notable adhesion virulence factors include intimin which is encoded by the *eae* gene within the Locus of Enterocyte Effacement (LEE), responsible for attaching-effacement lesions (13), and the aggregative adherence fimbriae which mediate adhesion between the bacteria and host cell in a “stacked-brick” profile (14). In 2018, the Food and Agriculture Organization of the United Nations (FAO)/World Health Organization (WHO) Joint Expert Meeting on Microbiological Risk Assessment (JEMRA) conceptualised a system (henceforth referred to as STEC JEMRA risk levels) to rank STEC according to their pathogenic potential into five STEC JEMRA risk levels, based on the *stx* subtypes and key adhesion molecules present (6). Building upon this concept, an analysis into clinical STEC within the European Union and European Economic Area also concluded that *stx* subtypes should also be considered during pathogenicity assessment (7). This report also noted that serotypes are not good indicators of severe clinical outcomes, and that while virulence factors for adhesion are a contributing factor, they are not essential for severe outcomes (7). Subtyping of *stx* can be performed by PCR (5), or inferred from whole genome sequencing (WGS) using tools like VirulenceFinder (15) and STECFinder (16).

Previously, we proposed a STEC virulence barcode to capture subtyping information of both *eae* and *stx* (17). This 12-mer barcode is divided into three segments: segment 1 (2-mer) encodes the *eae* subtype, segment 2 (2-mer) encodes predicted multiple copies of isogenic *stx* genes, and segment 3 (8-mer) captures up to four *stx* subtypes within a single genome. In addition to *eae* and *stx* subtyping, a key design consideration for the STEC virulence barcode was the ability to represent inferences on multiple isogenic *stx* copies (multiple copies of identical, or near identical *stx* genes, within a genome) from short-read sequencing data. This was performed by normalising the mean read depth of the detected *stx* operon against the mean read depth of *recA*, a single copy gene marker (17). Tracking of this is beneficial for genomic surveillance as harbouring multiple isogenic *stx* copies has been associated with increased disease severity (18, 19). Computing this virulence barcode relied upon multiple manual curation steps and multiple database searches (17).

Here we present a software tool to automatically generate the <u>STEC</u> virulence barc<u>ode</u> named “STECode”. We described changes to the original workflow by updating and adapting databases from other published sources. In addition, we also applied STECode to interrogate the virulence landscape of human STEC isolated from clinical samples collected in the state of New South Wales (NSW), Australia during the international border closure related to COVID-19 from April 2020 to February 2022. STECode is available at https://github.com/CIDM-PH/Stecode.

## Materials and Methods

### Curation of *stx* subtyping databases stecodeDB and stecodeCON

The database of STECfinder version 1.1.0 was used a foundation for the *stx* subtyping database (stecodeDB) used by STECode. Additional published *stx* subtypes not captured by STECfinder, along with updates to subtype designation, were also incorporated (Table 1). The updated stecodeDB consisted of 153 sequences (Figure 1) which spanned four *stx*_1_ subtypes (*stx*_1a,_ *stx*_1c,_ *stx*_1d,_ *stx*_1e_) and 15 *stx*_2_ subtypes (*stx*_2a_ through *stx*_2o_). Checks were performed against a validation set of genomes to ensure that the utilisation of stecodeDB, in conjunction with ABRicate version 1.0.0 (https://github.com/tseemann/abricate), revealed that the correct *stx* subtype could be recalled from genomic assemblies if > 21% of the stx operon could be assembled from short read sequencing (Supplementary Material). Intra *stx* nucleotide identities ranged from 96.87 % to 100 % while inter *stx* nucleotide identities ranged from 70.96 % to 99.76 % (Supplementary Material).

**Figure 1:**
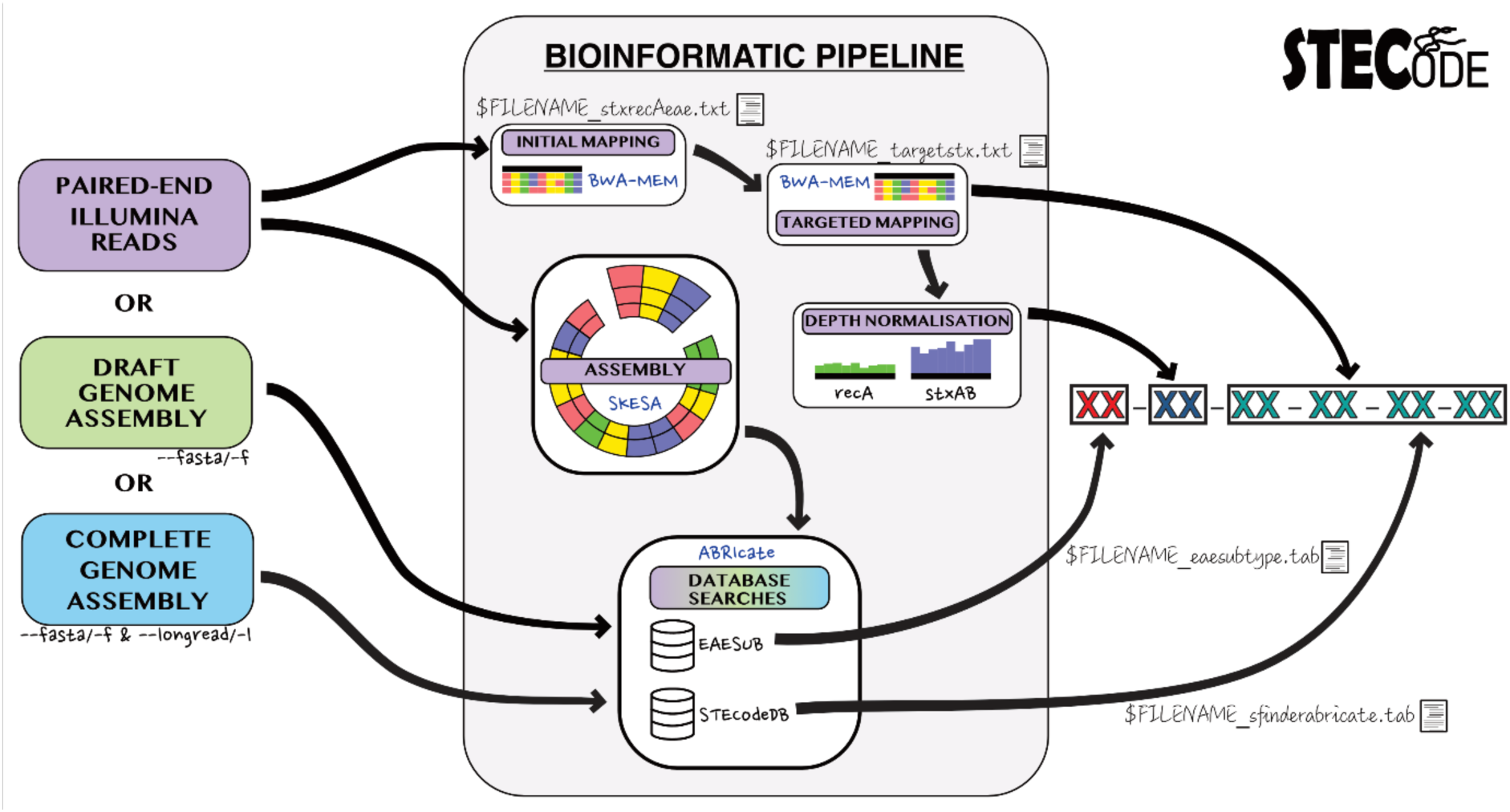
Distribution of the 153 *stx* subtype sequences within stecodeDB. For the design of STECode, the prototypical Shiga Toxin (*stx*) sequences from *Shigella* species were consolidated within *stx*_1a_ Image generated using ggplot version 3.4.2. (22).

**Table 1:** Modifications made to the STECFinder database for use in STECode.

| Database ID<sup>[a]</sup> | Reason for database modification | Reference |
| --- | --- | --- |
| **Removed from stecodeDB** |  |  |
| AB071620 (*stx*<sub>1c</sub>) | Original submission did not capture intergenic region between the A and B subunits. | (23) |
| AB071622 (*stx*<sub>1c</sub>) | Original submission did not capture intergenic region between the A and B subunits. | (23) |
| AB071624 (*stx*<sub>1c</sub>) | Original submission did not capture intergenic region between the A and B subunits. | (23) |
| **Reclassified for stecodeDB** |  |  |
| AM904726 (*stx*<sub>2l</sub>) | Reclassified from *stx*<sub>2e</sub> to *stx*<sub>2l</sub> as previously noted. | (7, 24) |
| **Sequences trimmed** |  |  |
| AM939642 (*stx*<sub>2e</sub>) | Trimmed to end at the stop codon of B subunit. | - |
| AY332411 (*stx*<sub>2e</sub>) | Trimmed to end at the stop codon of B subunit. | - |
| AJ567998 (*stx*<sub>2e</sub>) | Trimmed to end at the stop codon of B subunit. | - |
| **Added to stecodeDB** |  |  |
| KF926684 (*stx*<sub>1e</sub>) | Not currently in STECFinder version 1.1.0 | (7, 8) |
| MZ571121 (*stx*<sub>2j</sub>) | Not currently in STECFinder version 1.1.0 | (7) |
| KC339670 (*stx*<sub>2k</sub>) | Not currently in STECFinder version 1.1.0 | (7) |
| ERR4297034 (*stx*<sub>2m</sub>)<sup>[b]</sup> | Not currently in STECFinder version 1.1.0 | (9) |
| ERR4297035 (*stx*<sub>2m</sub>)<sup>[b]</sup> | Not currently in STECFinder version 1.1.0 | (9) |
| OQ054797 (*stx*<sub>2m</sub>) | Not currently in STECFinder version 1.1.0 | (9) |
| CP113092 (*stx*<sub>2n</sub>) | Not currently in STECFinder version 1.1.0 | (11) |
| CP113094 (*stx*<sub>2n</sub>) | Not currently in STECFinder version 1.1.0 | (11) |
| MZ229604 (*stx*<sub>2o</sub>) | Not currently in STECFinder version 1.1.0 | (10) |

[a]: *stx* subtypes listed in parenthesis.
[b]: Single-end reads were assembled using SPAdes version 3.15.5 (25) with the “--iontorrent” flag. Assemblies were interrogated using Artemis version 18.2.0 (26) and nucleotide sequences spanning the A and B subunits were extracted.

A second database was also generated (stecodeCON), which contained a single consensus sequence of each *stx* subtype to minimise the impact of intra and inter subtype cross mapping when paired-end Illumina data was used as an input. Intra-subtype multi-alignment of FASTA sequences (for subtypes with greater than one sequence) was performed using MAFFT version 7.511 (20) with the “--auto” flag and consensus sequences were generated from the multi sequence alignment using the em_cons program implemented within EMBOSS version 6.6.0.0 (21).

### Genomes and short reads for STECode validation

To validate the STECode pipeline, we consolidated a validation set of 25 publicly available diverse STEC and three non-*stx* encoding *E. coli* genomes (Table 2). Each selected STEC genome harboured between one and four *stx* operons, spread across different *stx* subtypes. To validate the ability to pick up isogenic *stx* copies by STECode, corresponding short reads of the 25 STEC genomes were also obtained (Table 2). Subtypes of *stx* and *eae* that were not reported within their corresponding publication were determined using STECfinder version 1.1.0 (16), and a custom *eae* subtype database respectively, and assumed as truth. Short reads from the three non-STEC control genomes were simulated using InSilicoSeq version 1.5.4 (27) to yield 1.5 million reads based on the MiSeq (Illumina) instrument.

**Table 2:** List of the 28 genomes used for the development of STECode.

| Genome[a] | Accession numbers |  | *stx* content | *eae* subtype<sup>[b]</sup> | Source |
| --- | --- | --- | --- | --- | --- |
|  | Chromosome | Short-reads |  |  |  |
| 194195 | CP044350.1 | SRR5024219 | *stx*<sub>2a</sub>, *stx*<sub>2c</sub><sup>[c]</sup> | γ1 (06) | (28) |
| 95JB1* | CP021335.1 | SRR954273 | *stx*<sub>1a</sub>, *stx*<sub>2a</sub> | γ2/θ (07) | (19) |
| 95NR1*# | CP021339.1 | SRR953500 | *stx*<sub>1a</sub>, *stx*<sub>2a</sub>, *stx*<sub>2a</sub>, *stx*<sub>2a</sub> | γ2/θ (07) | (19) |
| AUSMDU00002545* | CP045975.1 | SRR10520171 | *stx*<sub>1a</sub>, *stx*<sub>2c</sub><sup>[d]</sup> | γ1 (06) | (29) |
| AUSMDU00014361* | CP045827.1 | SRR8625824 | *stx*<sub>2a</sub> | β1 (03) | (29) |
| FWSEC0001 | CP031922.1 | SRR7896252 | *stx*<sub>1a</sub>, *stx*<sub>2a</sub> | β1 (03) | (30) |
| FWSEC0002 | CP031919.1 | SRR7896251 | *stx*<sub>1a</sub> | γ1 (06) | (30) |
| FWSEC0003 | CP031916.1 | SRR7896250 | *stx*<sub>1a</sub> | ε1(08) | (30) |
| FWSEC0004 | CP031913.1 | SRR7896257 | *stx*<sub>1a</sub>, *stx*<sub>2a</sub> | γ1 (06) | (30) |
| FWSEC0005 | CP031912.1 | SRR7896256 | *stx*<sub>1a</sub>, *stx*<sub>2a</sub> | γ2/θ (07) | (30) |
| FWSEC0006 | CP031910.1 | SRR7896255 | *stx*<sub>2a</sub> | ε1(08) | (30) |
| FWSEC0007 | CP031908.1 | SRR7896254 | *stx*<sub>1a</sub> | ε1(08) | (30) |
| FWSEC0008 | CP031906.1 | SRR7896259 | *stx*<sub>2a</sub>, *stx*<sub>2d</sub>, *stx*<sub>2d</sub> | Absent (00) | (30) |
| FWSEC0009 | CP031902.1 | SRR7896258 | *stx*<sub>2a</sub>, | Absent (00) | (30) |
| FWSEC0010 | CP031898.1 | SRR7896248 | *stx*<sub>2a</sub>, *stx*<sub>2d</sub> | Absent (00) | (30) |
| MBT-5 | CP058682.2 | SRR12207194 | *stx*<sub>1a</sub>, *stx*<sub>2a</sub> | β1 (03) | (31) |
| P15-385# | CP075627.1 | SRR13209798 | *stx*<sub>2a</sub>, *stx*<sub>2d</sub>, *stx*<sub>2d</sub> | Absent (00) | (32) |
| PNUSAE009425 | CP113092.1 | SRR6128619 | *stx*<sub>2n</sub> | Absent (00) | (11) |
| RM10466# | CP028381.1 | SRR7828259 | *stx*<sub>2a</sub>, *stx*<sub>2a</sub>, *stx*<sub>2d</sub> | Absent (00) | (33, 34) |
| STEC2017-197 | CP075663.1 | SRR13209817 | *stx*<sub>2d</sub>, *stx*<sub>2d</sub> | Absent (00) | (32) |
| STEC306 | CP091016.1 | SRR16071839 | *stx*<sub>2l</sub><sup>[e]</sup> | Absent (00) | (24, 35) |
| STEC307 | CP091018.1 | SRR16071838 | *stx*<sub>2l</sub><sup>[e]</sup> | Absent (00) | (24, 35) |
| STEC308 | CP091020.1 | SRR16071827 | *stx*<sub>2l</sub><sup>[e]</sup> | Absent (00) | (24, 35) |
| STEC865* | CP077649.1 | SRR15293247 | *stx*<sub>1c</sub> | Absent (00) | (36) |
| STEC866* | CP077646.1 | SRR15293246 | *stx*<sub>1c</sub> | Absent (00) | (36) |
| MG1655 | U00096.3 | Simulated reads | *stx* negative *E. coli* | Absent (00) | (37) |
| EC958 | HG941718.1 | Simulated reads | *stx* negative *E. coli* | Absent (00) | (38) |
| E2348/69 | FM180568.1 | Simulated reads | *stx* negative *E. coli* | α1 (01) | (39) |

### Bioinformatics tools used during STECode development

Read mapping of paired-end short reads were performed using BWA-MEM version 0.7.17 (40) using default settings. Output files from short-read mapping were further parsed using Samtools version 1.16.1 (41). Mapping results were visualised on the Integrative Genomics Viewer version 2.16.1 (42). Extraction of mapped reads from BAM files and conversion into FASTQ files were performed using SAMtools version 1.16.1 (41). Unless explicitly mentioned, assembly of short reads were performed using SKESA version 2.4.0 (43) using default settings. Gene detection via BLASTN in this study was performed using ABRicate version 1.0.0 (https://github.com/tseemann/abricate). Packaged ABRicate databases used in this study were EcOH (44) and VFDB (45). Custom ABRicate databases for this study were created in accordance with instructions from the GitHub page. Unless otherwise stated, default settings for ABRicate were utilised. Where required, assemblies were visualised using Artemis version 18.2.0 (26) and pairwise BLASTN comparisons were visualised using either Artemis Comparison Tool version 18.2.0 (46) or Easyfig version 2.2.2 (47). STECfinder version 1.1.0 (16) and VirulenceFinder 2.0 (15) were utilised to infer *stx* subtypes using default settings. VirulenceFinder 2.0 (*stx* database commit: 2023-07-20) was run locally as opposed to uploading sequences onto the web server.

### Overview of STECode pipeline

STECode was constructed as a pipeline written in Python to generate the three-part STEC virulence barcode (Figure 2). The pipeline accepts either paired-end short reads in FASTQ format or genomic assemblies in FASTA format. For assemblies, users can supply either draft assemblies (--fasta/-f flag) or completed assemblies (--longread/-1 flag). Declaration of these flags will bypass both genomic assembly and detection of multiple isogenic *stx* copies from short reads (Figure 2). In these cases, the segment 2 of the barcode is set to either “DG” or “CG” for draft or completed assemblies, respectively.

**Figure 2:**
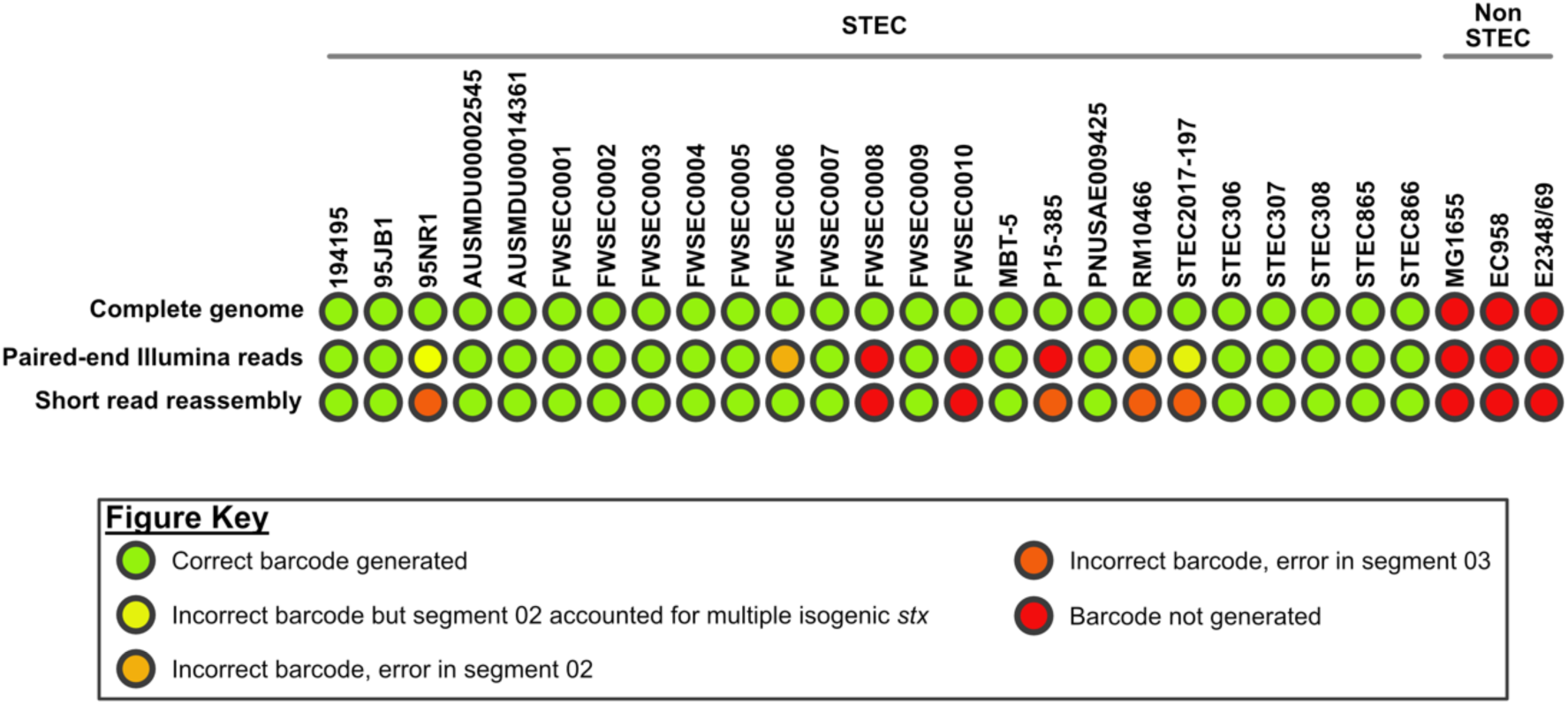
Graphical representation of STECode workflow. Pathways for the generation of the STEC virulence barcode from the three different inputs are shown. Key interim files generated by STECode useful for end user trouble shooting are also listed in the figure.

The standard STECode pipeline begins with the mapping of FASTQ reads to the stxrecaeae database (See supplementary material for database contents). Once mapped, all sequences with coverage greater than 98% are extracted and used as a reference sequence for a subsequent round of mapping (Targeted mapping). The normalisation value, derived from targeted mapping, is calculated, and used to infer the possibility of multiple, isogenic copies of the *stx* genes within the genome for segment 2 of the virulence barcode. Mapping results were tested on the short-read validation set to obtain the thresholds used by STECode (Supplementary Material). Possible codes for segment 2 include “00” (unlikely to possess multiple isogenic copies of *stx*), “T1” (possible multiple copies of a *stx*_1_ subtype), “T2” (possible multiple copies of a *stx*_2_ subtype),“01” (plausible multiple copies of a *stx*_1_ subtype), and “02” (plausible multiple copies of a *stx*_2_ subtype). After mapping, the supplied FASTQ is assembled with SKESA and subtypes of *eae* and *stx* (--mincov 21) are inferred using ABRicate against the eaesub and stecodeDB databases, respectively. Outputs of both *eae* subtyping and *stx* subtypes are used to populate segment 1 and segment 3 of the virulence barcode, respectively. Intermediate files and process logs are retained to aid in troubleshooting, in the instance of discrepant results between the mapping and assembly processes. STECode’s process considerations (thresholds & cut offs) are documented in the supplementary material.

### Culture conditions and sequencing of STEC isolated during the international border closure related to COVID-19

To examine the utility of STECode, we applied it to a set of STEC isolates that were collected in NSW as part of routine public health laboratory surveillance. STEC were isolated from clinical specimens by the Centre for Infectious Disease and Microbiology Laboratory Services (CIDMLS), Institute of Clinical Pathology and Medical Research (ICPMR), NSW Health Pathology, as per STEC control guidelines from Health Protection NSW (https://www.health.nsw.gov.au/Infectious/controlguideline/Pages/haemo.aspx). STEC studied (Supplementary Table S1) were isolated and processed for WGS on the Illumina NextSeq500 platform as previously described (17). All isolates were archived in Skim milk-Tryptone-Glucose-Glycerine (STGG) medium (48) at -80°C. When needed, isolates for long-read nanopore sequencing were recovered from -80°C, streaked onto Horse Blood Agar Columbia (HBA) plates (ThermoFisher Scientific), and incubated at 35°C for 16 ± 2 hours. A single colony from the overnight incubation was subsequently picked and subcultured onto a fresh HBA plates (ThermoFisher Scientific) and incubated at 35°C for 16 ± 2 hours. DNA extraction for long-read sequencing was performed as previously described (20) and libraries were prepared using the SQK-RBK004 kit (Oxford Nanopore Technologies) kit. Libraries were loaded onto a R9.4.1 flowcell and sequenced on a GridION running MinKNOW version 24.10. Passed (Q score > 9) FASTQ files were filtered using Filtlong version 0.2.1 (https://github.com/rrwick/Filtlong). Filtered long-reads were assembled using Flye version 2.9.2-b1786 (49) and long read polished using Medaka version 1.7.2 (https://github.com/nanoporetech/medaka). Confirmation of circularity was performed by visual inspection of the assembly graph file generated by Flye using Bandage version 0.8.1 (50). Annotation was performed using Prokka version 1.14.6 (51) running on metagenome mode (--metagenome) and genetic code 11. Insertion sites of *stx* prophages were determined via pairwise BLASTN alignment with the genome of *E. coli* K-12 strain MG1655 (GenBank accession: U00096.3), which was visualised using Artemis Comparison Tool (46). Insertion sites were further confirmed by BLASTN analysis of the phage borne. Phage genomes used for genomic comparison in this study include AU6Stx1 (GenBank accession: KU977420.1), Phage 2851 (GenBank accession: FM180578.1) (52) and Stx1a_EH2115 (GenBank accession: LC744180.1) (53).

## Results

### Accuracy of STECode barcode generation from the validation test set

Paired-end Illumina reads (simulated or published), short-read genomic reassembly and complete chromosome of the 28-validation set (25 STEC & 3 non-STEC) were each analysed on STECode to generate their respective STEC virulence barcode (Supplementary Table S2). When complete genomes were used as input, 25 STEC genomes generated their expected STEC virulence barcode while all three non-STEC complete genomes failed to generate the barcode (Figure 3). Regardless of the type of data input, all non-STEC failed to result in the generation of an STEC virulence barcode (Figure 3).

**Figure 3:**
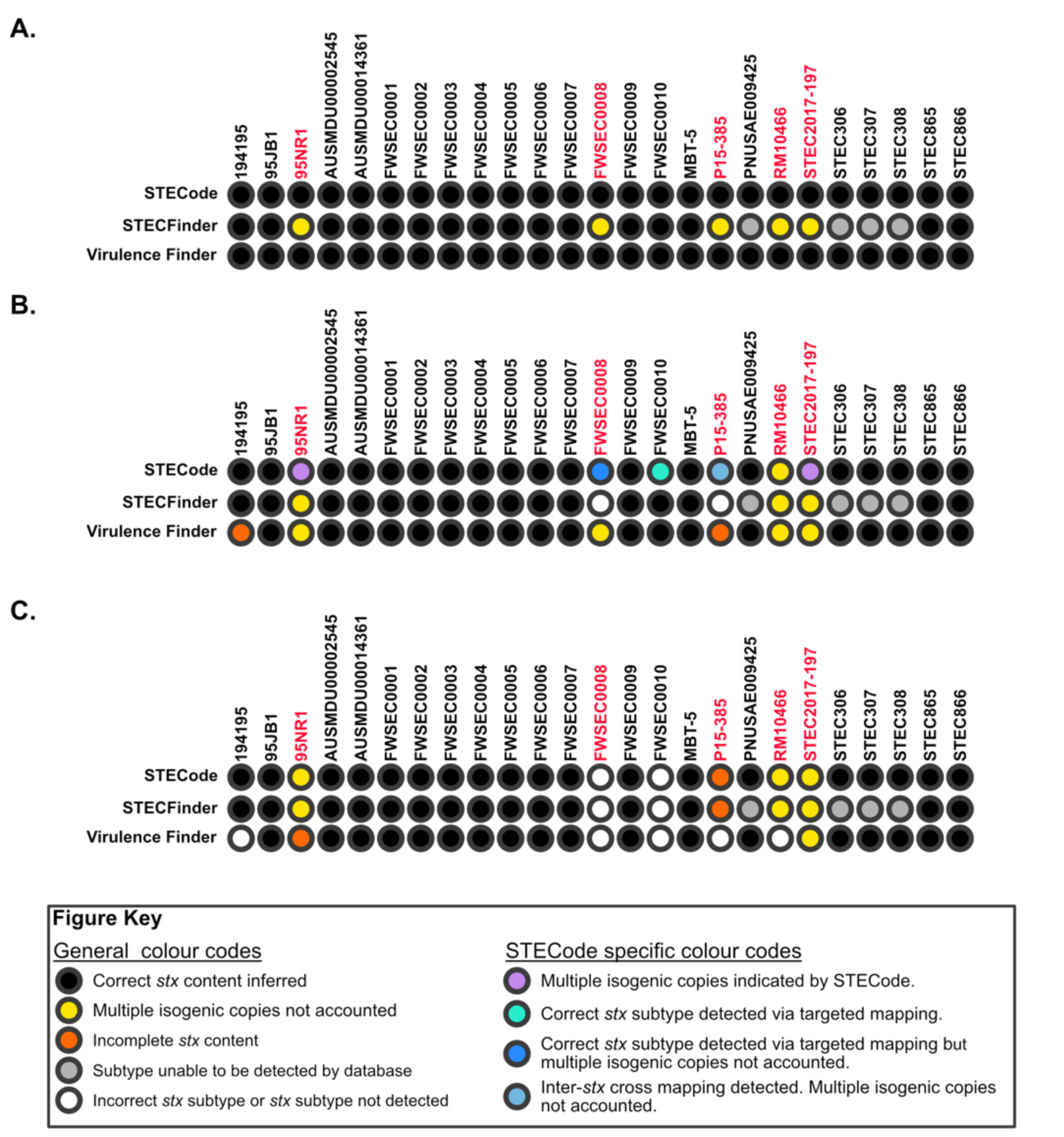
Graphical representation of concordance from STECode virulence barcode generation. Outcomes from STECode analysis on the validation set are colour-coded as per the figure key. Detailed results can be found in Supplementary Table S2.

When the paired-end Illumina reads were used as input, STEC virulence barcodes were generated from 22 out of 25 STEC genomes (Figure 3). Of these 22 STEC virulence barcodes, four genomes resulted in virulence barcodes that were discrepant with their respective *stx* content. The genomes of 95NR1 and STEC2017-197 each had three copies and two copies of *stx*_2a_ and *stx*2d copies, respectively (Table 2) but the generated barcode did account for these multiple copies within segment 02 of the barcode (Figure 3; Supplementary Table S2). The barcodes of FWSEC0006 and RM10466 each had errors in segment 02 within their barcode. The normalisation value for FWSEC0006 (*stx*_2a_ only) was 2.02 and for RM10466 (*stx*_2a_, *stx*_2a_, *stx*_2d_), the normalisation values were 1.68 and 1.06 for *stx*_2a_ and *stx*_2d,_ respectively, and this affected the correct inferences for segment 02. The remaining three STEC that did not result in the generation of the virulence barcode were FWSEC0008, FWSEC0010, and P15-385. The *stx* operons of FWSEC0008 and FWSEC0010 did not result in significant assembly (> 21% of the *stx* operon) and this lack of assembly led to the non-generation of the barcode (See supplementary material). When the interim files (targeted mapping) were interrogated for both FWSEC0008 and FWSEC0010, the correct *stx* subtypes could be detected at the sequencing reads level. Non generation of the barcode in P15-385 was due to both cross-mapping (with *stx*_2c_) and non-contiguous assembly of the *stx*_2d_ operon. Interim files (targeted mapping) also showed the presence of both *stx*_2a_ and *stx*_2d_, indicating that the correct *stx* subtypes could be detected in the sequencing reads level. Our validation set also included five STEC (Table 1) with insertion sequence (IS) element related disruptions to their *stx* operon, albeit all five resulted in a barcode (Supplementary Table S1). When the interim files of these five STEC were interrogated, traces of truncations were detected in assembly subprocess files ($Filename_sfinderabricate.tab) from four STEC, within the ‘coverage’ column, showing either missing bases (AUSMDU00002545) or as multiple entries with overlapping positions (STEC306, STEC307, STEC308). Small truncation could not be detected in 194195 as the 5’ end of the stx_2c_ operon could not be assembled, likely due to sequence similarity in this region with the stx2a operon that it also harboured. If judged solely on *stx* content (defined as all *stx* operons detected within the genome; regardless of truncation status), running STECode on paired-end Illumina reads resulted in a correct generation of the virulence barcode in 72% of the validation set. However, if the ability to infer multiple, isogenic *stx* copies was considered as beneficial, a correct barcode was generated for 80% of the validation set.

The issues identified in the pair-end Illumina sequencing dataset also had a corresponding impact when their short-read reassemblies were used as input (Figure 3; Supplementary Table S2). The only dataset with no corresponding impact was FWSEC0006, where the high normalisation value of its *stx*_2a_ had no bearing on an assembly-based barcode generation method. Like when using paired-end Illumina reads as input, both FWSEC0008 and FWSEC0010 could not generate a barcode due to the non-assembly of their *stx* operons (See supplementary material). Despite having non-contiguous assembly of the *stx*_2d_ operon, the short-read reassembly of P15-385 could assemble its *stx*_2a_ operon and thus, an erroneous barcode was generated. Re-assemblies of 95NR1, STEC2017-197, and RM10466 each did not generate a barcode that captured its entire *stx* content, but this was expected as multiple, isogenic *stx* copies will collapse into a single contig when short-read assembled. When judged solely on *stx* content, the utilisation of short-read reassemblies on STECode resulted in the generation of the correct virulence barcode in 76% of the validation set.

### Comparison of STECode *stx* subtyping with VirulenceFinder and STECFinder

Both VirulenceFinder and STECfinder were run on the 25 STEC validation set genomes and compared to the output of STECode (Figure 4; Supplementary Table S3). With a completed genomic assembly as input, both STECode and VirulenceFinder identified the correct *stx* content. STECFinder, on the other hand, could not accurately report the *stx* content of STEC with multiple, isogenic *stx* copies with multiple copies collapsing into a single copy (Supplementary Table S3). In addition, the current version of STECFinder does not include sequences of *stx*_2l_ and *stx*_2n_ thus, the *stx* content of PNUSAE009424, STEC306, STEC307, and STEC308 could not be reported on.

**Figure 4:**
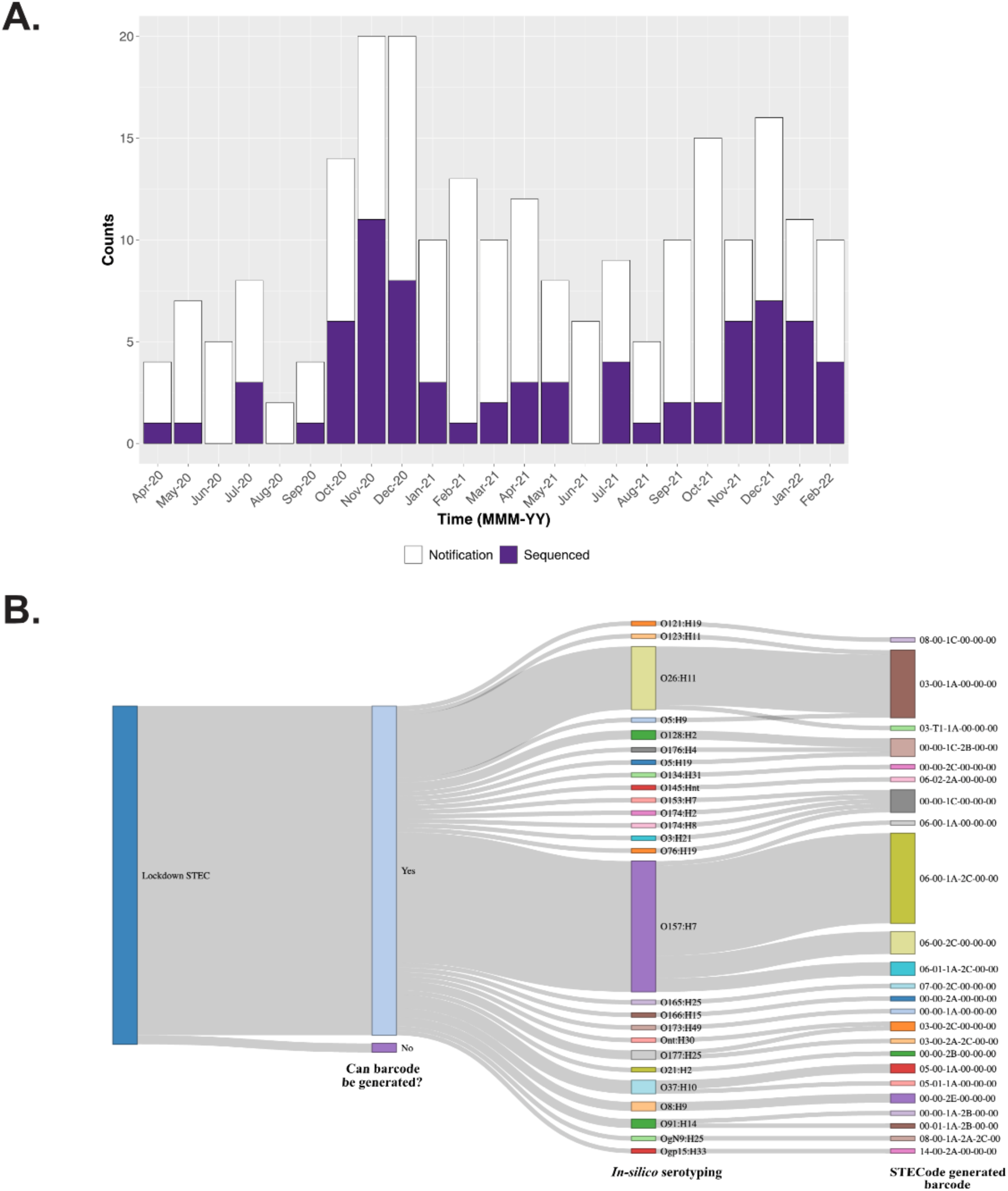
Graphical representation of concordance with *stx* content from STECode, STECFinder and VirulenceFinder. Inputs for the three software include (A) complete genomes, (B) pair-end Illumina reads, and (C) short read reassemblies. Genomes with multiple isogenic *stx* copies are labelled in red. Interpretations of *stx* content identification are colour-coded as per the figure key. Detailed results can be found in Supplementary Table S3.

When paired-end Illumina reads were used as input, the inability to directly report on the *stx* content from STEC genomes (n =5) with multiple, isogenic copies of *stx* were observed using all three software (Figure 4B). Only STECode was able to infer the presence of multiple, isogenic copies of *stx,* albeit with only two of the five validation STEC genomes. Inability to directly report on *stx* content in these five genomes was also observed when draft assemblies were used as input (Figure 4C). Overall, our results showed that while STECode, STECFinder, and VirulenceFinder are comparable in their ability to detect *stx* content, STECode has an additional advantage of detecting the presence of multiple, isogenic copies of *stx*.

### Pathogenic potential of circulating STEC within NSW during the COVID-19 border closure

During the COVID-19 pandemic, Australia closed its borders to all non-Australian travellers on the 15^th^ of March 2020 and reopened borders to fully vaccinated visa holders on the 21^st^ of February 2022. From April 2020 to February 2022, there were 229 STEC infections notified in NSW and of these, 75 resulted in successful culture and were sequenced using short-read sequencing on the Illumina platform (Figure 5A). The 75 sequenced STEC were split across 27 serotypes (*in-silico* serotyping) with serotype O157:H7 being the predominant serotype (30/75, 40%) followed by serotype O26:H11 (14/75, 18.70%). Cumulatively, these two serotypes made up 58.7% of our dataset during this period. When each genome was subjected STECode analysis, 73 of 75 genomes yielded a virulence barcode (Figure 5B; Supplementary Table S4).

**Figure 5:**
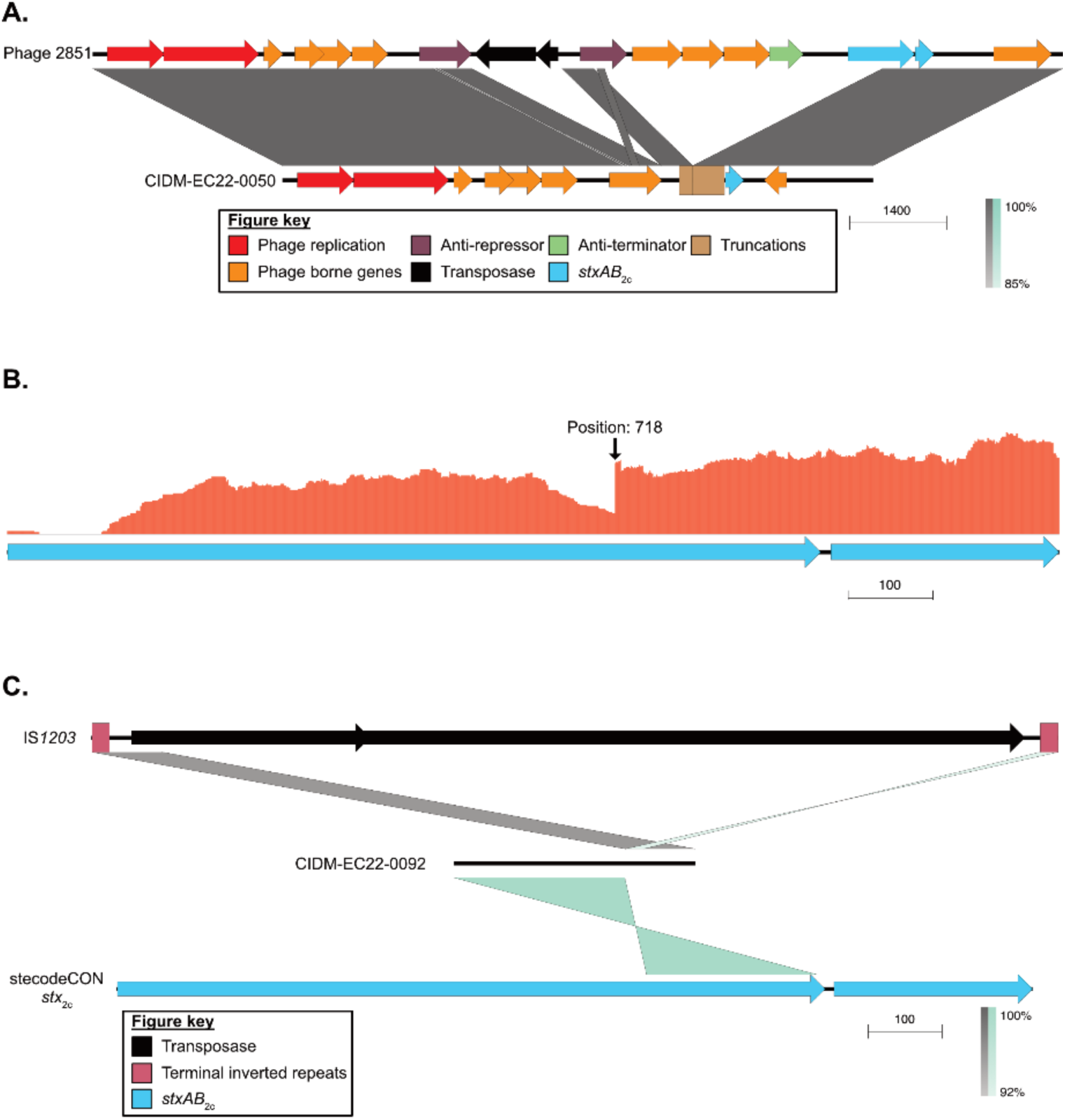
STEC sequenced during the COVID-19 lockdown in NSW. (A) Group barplot of STEC notifications in NSW and number of STEC sequenced each month during the lockdown period. Grouped barplot generated using ggplot2 version 3.4.2. (22). (B) Sankey diagram outlining the information flow from the 75 STEC sample, through to serotype and virulence barcode generation by STECcode. Sankey diagram was generated using the networkD3 package (https://CRAN.R-project.org/package=networkD3).

The most prominent STEC virulence barcode in our lockdown dataset was 06-00-1A-2C-00-00 (20/73, 27.40%), which corresponded to JEMRA Risk Level 3. This predominance was due to the large proportion of STEC O157:H7 present (Figure 5B). STEC belonging to JEMRA Risk Level 1 made up the minority (5/73, 6.84%) while strains representing JEMRA risk level 3 (32/75, 43.84%) and JEMRA risk level 4 (25/73, 34.25 %) represented the largest and the second largest proportion in our dataset, respectively (Figure 5B & Supplementary Table S4). As expected from a phage borne virulence factor, stratification of virulence barcodes with serotype was not observed. Genomes belonging to both STEC O157:H7 and STEC O177:H5 each had multiple barcodes within their respective cohort, reflecting on their diversity in *stx* content. Conversely, virulence barcodes like 00-00-1C-00-00-00 and 00-00-1C-2B-00-00 can be found in multiple STEC serotypes (Figure 5B).

### Truncation of *stx*2c detected in the NSW dataset

CIDM-EC22-0050 (STEC O157:H7) and CIDM-EC22-0092 (STEC O49:H16) did not yield a barcode due to each genome having a discrepant *stx*_2c_ result (Supplementary Table S4). When the mapping files ($Filename_stxrecAeae.tab) generated by STECode were interrogated, it was revealed that mapping across *stx*_2c_ yielded a query coverage of 59.47% and 94.12% for CIDM-EC22-0050 and CIDM-EC22-0092, respectively. There values were below the threshold set by STECode to proceed onto targeted mapping and thus *stx*_2c_ could not be detected. At the assembly level, the *stx*_2c_ operon of CIDM-EC22-0050 was assembled within an 8,549 bp contig with a truncation of 498 bases at the 5’ of *stxA*_2c_. Further interrogation revealed that the coding sequence upstream of the truncated *stxA*_2c_ was not the expected phage anti-terminator but a truncated anti-repressor (Figure 6A). Pair-wise nucleotide comparison against the genome of *stx*_2c_ phage 2851 (GenBank accession: FM180578) revealed a large-scale deletion event within the *stx*_2c_ prophage of CIDM-EC22-0050 (Figure 6A). The mechanism of deletion was not apparent but nonetheless, the truncation of *stxA*_2c_ indicated that CIDM-EC22-0050 would be unlikely to produce Stx2c.

**Figure 6:**
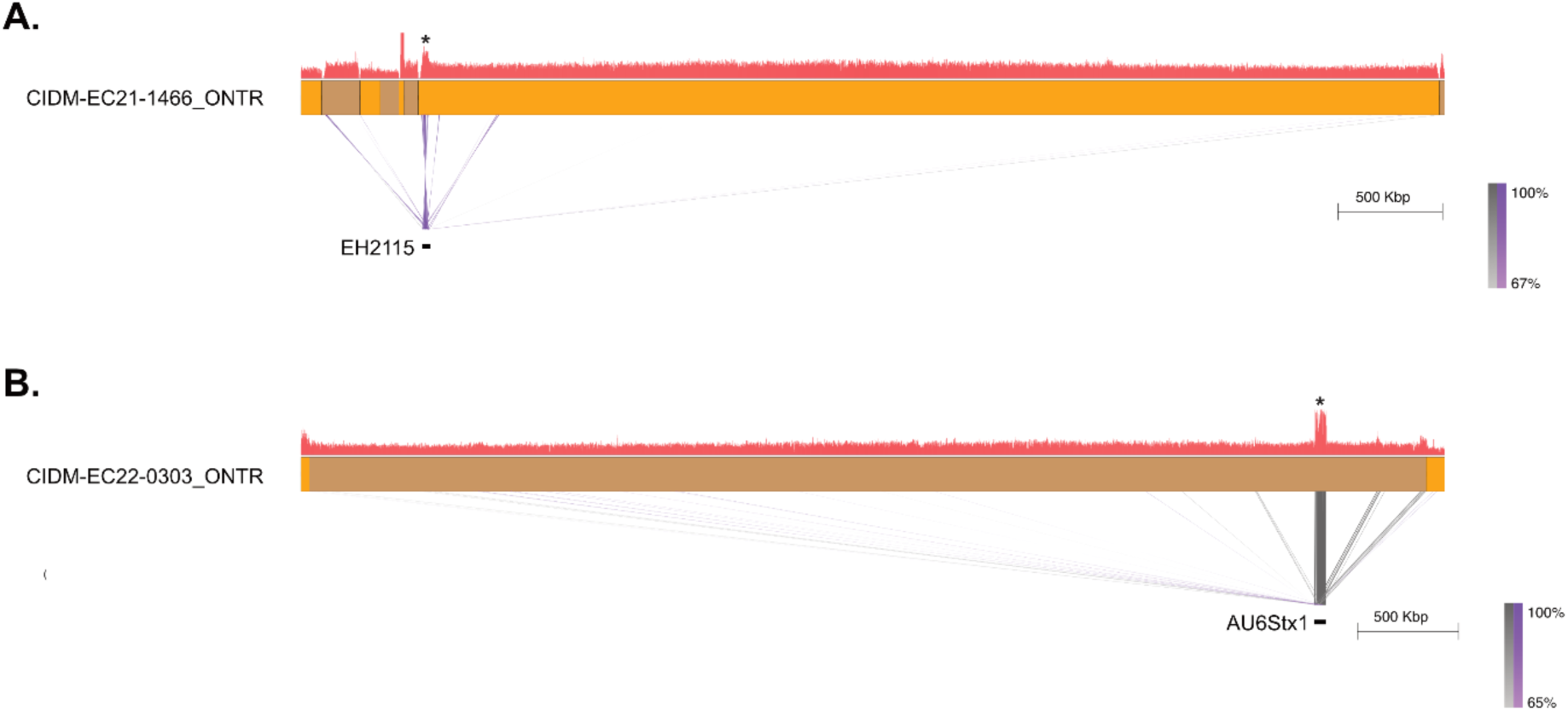
Truncations in *stxAB*_2c_ in both CIDM-EC22-0050 and CIDM-EC22-0092. **(A)** Pairwise BLASTN comparison between Phage 2851 (Top; GenBank accession: FM180578.1; genomic positions: 11,762 - 25,810 bp) against the *stx*_2c_ containing contig in CIDM-EC22-0050 (Bottom). Coding sequences are colour coded according to the figure key. **(B)** Mapping profile of CIDM-EC22-0092 across the *stx*_2c_ consensus sequence of stecodeCON. Coding sequence of *stxAB*_2c_ are represented as arrows. **(C)** Pairwise BLASTN comparison between IS*1203* (Top; GenBank accession: U06468.1; genomic positions: 42 to 1,352 bp), contig with partial assembly of *stx*_2c_ in CIDM-EC22-0092 (Middle) and stecodeCON *stx*_2c_ consensus sequence (Bottom). Coding sequences are colour coded according to the figure key. For panels **(A)** and **(C)**, grey shading between the genomes in both panels represent BLASTN matches on the same strand while green shading represent BLASTN matches on the opposite strand. All panel figures were generated using Easyfig2 (47) .

The *stx*_2c_ operon in CIDM-EC22-0092 was assembled within a 1,756 bp contig without the first 721 bases (out of 1,241 bases; 58.10%). The mapping profile of CIDM-EC22-0092 across stecodeCON *stx*_2c_ sequence revealed the presence of clipped reads in a majority of reads at position 718 (48/68 reads; Figure 6B). When CIDM-EC22-0092 reads that were mapped to stecodeCON’s *stx*_2c_ sequence were collated and reassembled, the assembly resulted in three contigs of sizes 574 bp, 276 bp, and 328 bp with the latter contig consisting of positions 718 to 949 (232/328 bases) of *stx*_2c_. The remaining 96 bases were determined to the be first 96 bases of the insertion sequence IS*1203* (Figure 6C). Based on both the mapping and the local reassembly of this fragment, the evidence indicated that the *stxA*_2c_ in CIDM-EC22-0092 was truncated by IS*1203* and that this isolate was unlikely to express Stx2c.

### Unintended prophage induction can impact upon isogenic *stx* probability call (segment 02)

Seven STEC genomes during the border closure period were inferred to either possibly (1/7) or plausibly (6/7) harbour multiple, isogenic *stx* copies within their genome (Supplementary Table S7). Of these seven STECs, CIDM-EC20-1974 (00-01-1A-2B-00-00; normalisation value: 2.94), CIDM-EC21-1466 (05-01-1A-00-00-00; normalisation value: 2.18), and CIDM-EC22-0303 (06-01-1A-2C-00-00; normalisation value: 3.68) were selected for long-read sequencing (henceforth suffixed with _ONTR) to determine if they do indeed carry multiple, isogenic copies of *stx_1a_*.

Long-read sequencing data of these three STEC were filtered to keep all reads greater than 5,999 bases to aid in genomic reconstruction of the bacteria chromosome (Supplementary Table S5). The chromosome of both CIDM-EC20-1974_ONTR and CIDM-EC22-0303_ONTR each resolved into a single circular contig when the higher threshold was used (Supplementary Table S5). The chromosome of CIDM-EC21-1466 remained fragmented, even at the higher threshold, with a contig break at the tail morphogenesis region of the *stx*_1a_ prophage. When subjected to STECode analysis, barcodes of 00-CG-1A-1A-2B-00, 05-CG-1A-00-00-00 and 06-CG-1A-2C-00-00 were generated for CIDM-EC20-1974_ONTR, CIDM-EC21-1466_ONTR and CIDM-EC22-0303_ONTR, respectively. This indicated that only CIDM-EC20-1974 had additional copies of *stx*_1a_ with one *stx*_1a_ prophage integrated into *ynfG* (also known as *dmsB*) and another *stx*_1a_ prophage integrated into *wrbA*.

To determine if the plausibility scores of both CIDM-EC21-1466 and CIDM-EC22-0303 were inflated due to the capture of *stx* prophage induction, short-reads (from the initial DNA extraction) were mapped onto their corresponding long-read assemblies. Mapping profiles for both CIDM-EC21-1466_ONTR and CIDM-EC22-0303_ONTR showed elevated read coverage over their *stx*_1a_ prophages which were inserted into *torS*-*torT* and tRNA-*argW* respectively (Figure 7; Supplementary Table S5). This indicated that these *stx* prophages were induced either during the cultivation or the extraction process from these two STEC. Taken together, our results showed that only one STEC isolated during the COVID-19 lockdown period harboured multiple, isogenic *stx* copies.

**Figure 7:**
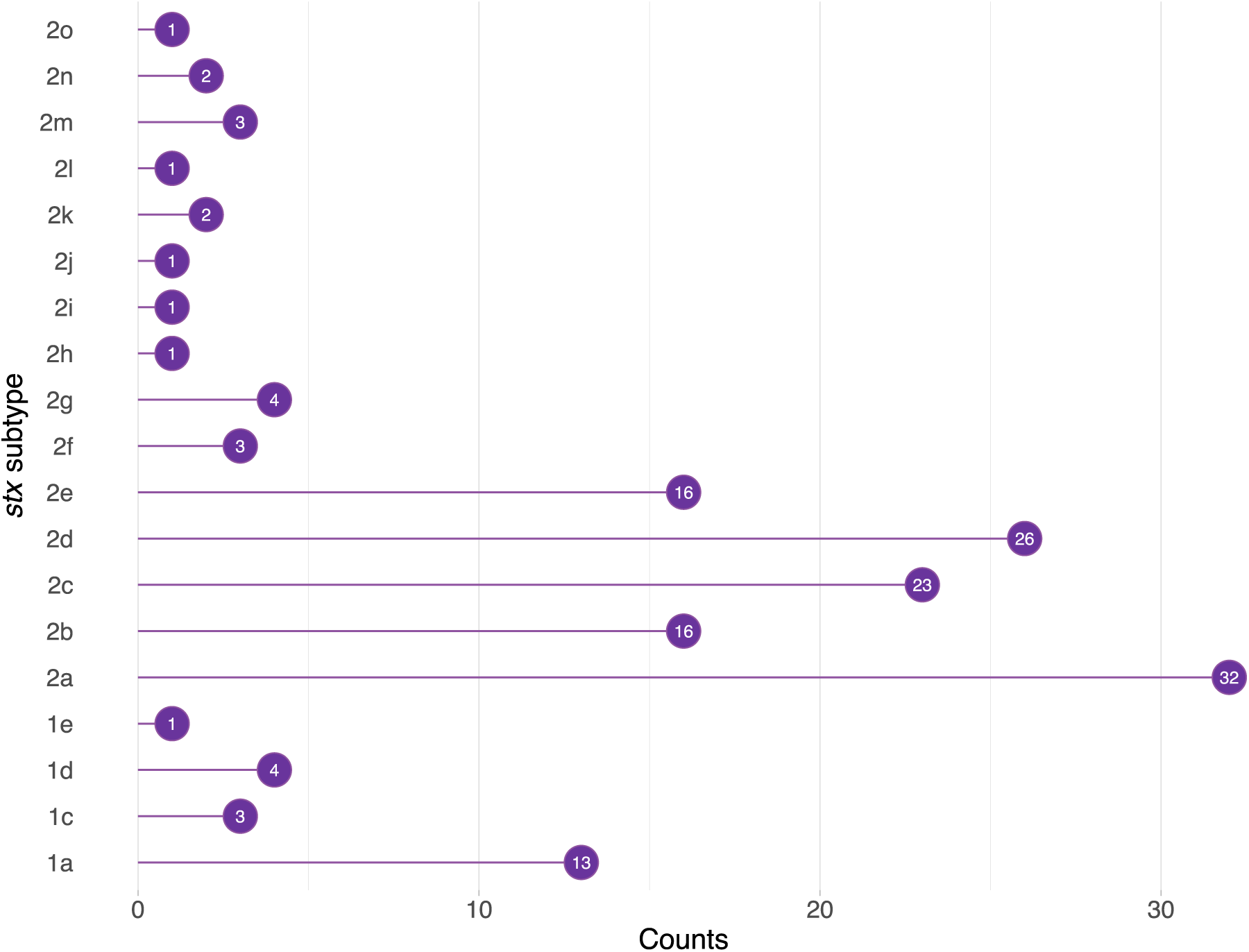
Induced prophages captured in paired-end Illumina data. **(A)** Pairwise BLASTN comparison between the long-read assembly of CIDM-EC21-1466 (Top) against Stx1a_EH2115 (Bottom; NCBI GenBank accession: LC744180.1). Contigs are colour coded in alternating orange and olive. Short read mapping profile over the assembly is represented as a histogram above the assembly with read depth as the Y-axis (Max-Y capped at 150x). Asterisk (*) denotes the position of the *stx*_1a_ prophage in CIDM-EC21-1466. **(B)** Pairwise BLASTN comparison between the long-read assembly of CIDM-EC22-0303 (Top) against the AU6Stx1 (Bottom; GenBank accession: KU977420.1). Short read mapping profile over the assembly is represented as a histogram above the assembly with read depth as the Y-axis (Max-Y: 300). Asterisk (*) denotes the position of the *stx*_1a_ prophage in CIDM-EC21-0303. In both panels, contigs are colour coded in alternating orange and olive and grey shading between the genomes represent BLASTN matches on the same strand while purple shading represent BLASTN matches on the opposite strand. All panel figures were generated using Easyfig2 (49).

## Discussion

Risk assessment for STEC is largely based upon the detection of *stx* subtypes present within its genome. This manuscript describes and extends considerations (See supplementary material) for the development and validation of STECode, a software tool to automatically generate the STEC virulence barcode to aid in genomic epidemiology. The STEC virulence barcode was initially conceptualised to meet the need of genomic surveillance and risk assessment of virulence, providing a quick snapshot of pertinent virulence factors to inform public health assessments. The STEC virulence barcode was not designed to be a standalone entity, but to be utilised in conjunction with other data points like serotyping, multi-locus sequence typing or core genome sequence typing, to support genomic surveillance activities.

### Different software, different “value-add” for genomic surveillance

At the base level, STECode, STECFinder, (16) and VirulenceFinder (15) can each perform *stx* subtyping from both sequencing reads and assemblies. However, they each differ in the type of “value-add” in terms of genomic surveillance. VirulenceFinder has a suite of additional databases to interrogate the virulome of the STEC genome of interest, beyond stx typing. STECFinder reports on serotype of the STEC in question, a classical epidemiological marker used for public health surveillance. A prerequisite requirement for STECode is the software ABRicate, which comes pre-packaged with databases useful for STEC analysis like EcOH (44) for *in-silico* serotyping and VFDB (45), or ecoli_vf (https://github.com/phac-nml/ecoli_vf) for virulence gene detection. However, the key focus of STECode is on identifying a blind spot from short-read WGS of STEC which was the ability to carry multiple copies of the same *stx* subtype in its genome (18) . The ability for *stx* phages to circumvent superinfection immunity (54), and STEC harbouring multiple copies of isogenic *stx* genes leading to severe clinical complications (18, 19) have both been described. It is therefore prudent for genomic surveillance of STEC to have an oversight of this occurrence.

While the possibility/plausibility of multiple isogenic *stx* copies can be flagged by STECode, it is by no means confirmatory. Previously we have acknowledged that artefacts from sample processing could impact upon inferences of extra *stx* copies (17). In this study, our findings provided circumstantial evidence for this occurrence with the three long-read sequenced STEC. Short-read sequencing showed that each genome plausibly harboured extra copies of the *stx*_1a_ gene, but subsequent long-read sequencing (not on the same extract) showed that this was true for only one STEC. Induced prophages in bacterial genomes can be detected via WGS and this event is represented by elevated read depth over their prophage regions in relation to its flanking chromosome genomic regions (55, 56). The two STEC genomes, which only had one copy of *stx*_1a_, had elevated read coverage across their respective *stx*_1a_ prophages. This indicated that the *stx*_1a_ prophage of both these STEC were induced at some point during culturing to DNA extraction. However, we did not investigate possible reasons for induction in both these STEC. Nonetheless, for genomes showing either possibility or plausibility of multiple isogenic *stx* copies, long-read sequencing (if available) for confirmation would be a prudent approach.

### STECode supports genomic context investigation and verification

One consideration for STECode is the providence of interim files to allow end-users to cross check their results. In both our validation and border closure dataset, there are genomes for which STECode did not generate a virulence barcode due to discrepancies detected in the output between the different sub-processes. Interrogation of these files helped guide the identification of structural variation within the *stx*_2c_ operon of CIDM-EC22-0050 and CIDM-EC22-0092, which would have led to the abolishment of toxin expression in both cases.

We would also like to encourage end-users to check intermediate files even when STECode generates a virulence barcode. The thresholds within STECode are set rather permissively for the detection of *stx* subtypes and consequently small truncation of the *stx* operon by either by IS elements (57–60) or other means can still result in barcode generation. Disruptions that occur within the *stx* operon are more likely to be picked up by the assembly subprocesses and can be interrogated from the provided interim files. However, small truncations at the 5’ or 3’ end are more likely to be missed. Depending on the complexity of the *stx* content, these truncations at either end could either be picked-up by the assembly subprocesses (e.g. in the validation genome AUSMDU00002545) or be completely masked due to assembly issues (like in the validation genome 19145).

### Utilisation of JEMRA risk levels

STECode was designed to report on the presence of *stx* subtypes within STEC genomes and to also facilitate inferences of STEC JEMRA risk levels. The JEMRA risk levels are dependent on the presence of key *stx* subtypes for attribution, an easy process for automation. However, attribution to JEMRA risk level was not considered for automation with a key consideration being disruptions from within (i.e. in the *stx* operon) or around (i.e. key modules of the *stx* prophage). Current evidence suggests that truncations to the *stx* genes can either disrupt toxin production (60) or have no noticeable effects (58). If Stx expression is disrupted (something that genomics cannot predict), it should alter the pathogenic potential of STEC. A case in point would be CIDM-EC22-0050 in this study which had an intact *stx*_1a_ but a truncated *stx*_2c_. This disruption meant that CIDM-EC22-0050 should be classified as a JEMRA risk level 4 instead of a JEMRA risk level 3. It is because of this potential misclassification that we took the conservative approach to not automatically assign JEMRA risk levels, leaving this association to the end-user after manual curation of the output files from STECode.

### Use with other Stx-producing bacteria

Genes encoding *stx* are phage borne, and *stx* have been previously detected in *Acinetobacter haemolyticus* (61), *Aeromonas caviae* (62), *Citrobacter freundii* (63, 64), *Enterobacter cloacae* (8), *Escherichia albertii* (65, 66), *Shigella flexneri* (67) and *Shigella sonnei* (68). While STECode was designed for use in STEC analysis, it has limited utility for other Stx producing organisms. A virulence barcode will not be generated if no reads map onto an *E. coli recA* gene but there, an interim file would nonetheless be produced with the *stx* subtyping result. The database behind STECode, which was built upon the *stx* database of STECFinder (16), also contained subtype sequences from other *stx* containing organisms and thus known subtypes would be captured. In the event of a suspected novel *stx* subtype in either STEC or other *stx* containing organisms, users can also leverage the intermediate files. In this study, we showed that inter *stx* subtype nucleotide percentage identity, within our STECode databases ranged between 70.96% to 96.66%. Using this cut-off as a guide, users can peruse the provided interim output files to assess the nucleotide percentage ID of the detected *stx* subtype within their respective Stx producing organism. If the best hit from STECode is within or below this range, this could serve as a signal to perform further analysis such as nucleotide and amino acid sequence alignments to confirm the novelty of the detected *stx* subtype (5, 8, 9, 69).

### NSW lockdown dataset indicated that STEC of lower pathogenic potential remain endemic

Prior to the COVID-19 border closures, it was reported that STEC isolated from clinical sources in NSW predominantly harboured *stx* genes of lower pathogenic potential (17). The COVID-19 lockdown provided an opportunity to interrogate the pathogenic potential STEC landscape without external influences. We showed that within our dataset, STEC of lower pathogenic potential were in circulation. This observation is consistent with the trend previously observed at the pre-lockdown period, both at the national or the state level, which showed that most clinical STEC had lower pathogenic potential (17, 70–75).

A key caveat on our observation is that the available STEC genomes do not necessarily reflect the STEC populations in circulation. The process of WGS is highly reliant upon the ability to culture the organism and in our study period, we only captured 32.80% of total notifications. During this same period, there was only one HUS notification (Obtained from the National Notifiable Diseases Surveillance System: https://nindss.health.gov.au/pbi-dashboard/) for the state of NSW. While we do not have genome sequences for more than 70% of the STEC, the lack of notified HUS cases provided indirect evidence that at least within the state of NSW, STEC of lower pathogenic potential are in circulation.

## Conclusion

In conclusion, STECode offers an accessible method for *stx* subtyping and the generation of the STEC virulence barcode for genomic surveillance as well as clinical and public health risk assessment. The monitoring of *stx* subtypes can add value to classical epidemiological tracking of STEC where serotype is a principal component. We envisaged that this tool is useful for genomic surveillance of STEC and inform genomic epidemiology investigation of this pathogen of public health concern.

## Author contributions

Eby M. Sim: Conceptualisation, Methodology, Validation, Investigation, Data Curation, Writing-Original draft, Writing - Review & Editing.

Winkie Fong: Conceptualisation, Software, Investigation, Data Curation, Writing-Original draft, Writing - Review & Editing.

Carl Suster: Software. Writing-Original draft, Writing - Review & Editing.

Jessica E. Agius: Visualisation, Writing - Review & Editing

Shona Chandra: Visualisation, Writing - Review & Editing Basel Suliman: Resources, Writing - Review & Editing.

Qinning Wang: Resources, Writing - Review & Editing.

Christine Ngo: Investigation, Writing - Review & Editing

Connor Finemore: Investigation, Writing - Review & Editing

Sharon C. A. Chen: Resources, Writing - Review & Editing

Kerri Basile: Resources, Writing - Review & Editing

Vitali Sintchenko: Conceptualisation, Resources, Supervision, Writing-Original draft, Writing

- Review & Editing.

## Acknowledgements

The authors would like to acknowledge staff both past and present of CIDMLS, ICPMR NSW Health Pathology. In particular, the Core Microbiology Unit, NSW Enteric Reference Laboratory, and the Microbial Genomics Reference Laboratory. The authors would also like to acknowledge the Sydney Informatics Hub along with The University of Sydney High Performance Computing Cluster for bioinformatics assistance and computing capacity.

## Data availability

Sequencing reads and assemblies generated in this study were uploaded to Bioproject: PRJNA1098252. Accession numbers for short read data can be found in Supplementary Table S1, while accession numbers for long-read fastq files can be found in Supplementary Table S5. Reassembled short-read assemblies of the 24-validation set along with the reassembly of the *stx*_2c_ in CIDM-EC22-0092 were uploaded to Figshare (10.6084/m9.figshare.24277228).

