## Supplementary Material for "STECode: an automated virulence barcode generator to aid clinical and public health risk assessment of Shiga toxin-producing *Escherichia coli*"

### Results

Results within this supplementary material details the design considerations for the development of STECode. For methods used in the supplementary material, please consult the "Bioinformatics software used during STECode development" section in the materials and methods of the main manuscript.

#### Establishment of truth: *stx* and *eae* subtyping of validation set

Of the selected 25 validation STEC set genomes, six did not have reported *stx* subtyping results and subtyping was performed on these genomes (See Table 2 of main manuscript). Running STECFinder using genomic assemblies as input would result in the collapse of multiple isogenic *stx* copies and as such, all complete genomes were manually curated. In addition to the three known STEC genomes with multiple isogenic *stx* copies, FWSEC0008 was also determined to harbour an additional copy of $stx_{2d}$. STEC306, STEC307, STEC308 each had a known IS*2* family insertion sequence inserted between $stxA_{2l}$ and $stxB_{2l}$, but manual curation also revealed small truncations in two other STEC genomes. The last 36 bases of $stxB_{2c}$ and the first 124 bases of $stxA_{2c}$ in AUSMDU00002545 and 194195 respectively were each truncated by an IS*1203* insertion. All *stx* subtypes detected in the validation-set were assumed as truth for the validation of STECode.

Unlike the subtyping of *stx*, subtyping results of *eae*, are rarely reported in the literature. To determine the presence of the LEE and subtypes of *eae*, complete genomes of the 24-validation set were subjected to BLASTN analysis against the VFDB and a custom ABRicate *eae* database (henceforth referred to as eaesub). Presence of *eae* and the LEE were detected 13 out of 24 genomes (Figure SM1). All LEE positive genomes were each assigned a virulence barcode *eae* designation as previously described (1). All *eae* subtypes detected in the validation set were assumed as truth for the validation of STECode.

#### Database adaptation for *stx* subtyping by STECode

The initial database (henceforth referred to as VB01) used for *stx* subtype inference was an adapted reference panel which consisted of 10 representative sequences spanning subtypes $stx_{1a}$, $stx_{1c}$, $stx_{1d}$, and $stx_{2a}$ through $stx_{2g}$ (1, 2). The database behind STECfinder version 1.1.0 consisted of 147 sequences (3), inclusive of the 10 representative sequences from VB01, and spanned subtypes $stx_{1a}$, $stx_{1c}$, $stx_{1d}$, $stx_{2a}$ through $stx_{2i}$ and $stx_{2k}$. Both databases cumulatively lacked sequences belonging to subtypes $stx_{1e}$, $stx_{2j}$, $stx_{2k}$, $stx_{2m}$ and $stx_{2}$. To capture these subtypes, along with subtype updates according to the literature, the database of STECfinder was used as a foundation and modified (See Table 1 of main manuscript).

Intra-subtype BLASTN comparison within stecodeDB had nucleotide identities ranging from 96.87% to 100% while inter-subtype BLASTN comparisons showed nucleotide identities ranging from 70.96 % to 99.76 % (Table SM1). The higher ranges within the inter-subtype comparison were due to high nucleotide similarities between the subtypes $stx_{2a}$, $stx_{2c}$ and $stx_{2d}$, which are known to be highly genomically similar (2). When $stx_{2a}$, $stx_{2c}$ and $stx_{2d}$ BLASTN hits were omitted from their respective inter-*stx* subtype comparison, the maximum nucleotide identity was reduced to ranges comparable with other inter-*stx* subtype comparisons (Table SM1).

When the equivalent subtype sequences of stecodeDB ($stx_{1a}$, $stx_{1c}$, $stx_{1d}$, and $stx_{2a}$ through $stx_{2g}$) were tested against VB01, concordance was observed in majority of the subtypes (Figure SM2). Discordance was observed amongst the $stx_{2d}$ sequences where majority of the $stx_{2d}$ sequences (18/26; 69.23%) in stecodeDB resulted in miscalls (Figure SM2). These 18 sequences were typed as either $stx_{2a}$ (2/18; 11.11%) or $stx_{2c}$ (16/18; 88.89%). Eight subtypes (11 sequences) in stecodeDB had no sequence representation in VB01. When these 11 sequences were tested against VB01, the misassigned nucleotide identities ranged from 86.97% to 96.13%, values within the range of misclassification amongst inter-*stx* subtypes (Tables SM1 & SM2).

#### Multiple "decoy" sequences not inducive for normalisation determination

Utilisation of stecodeDB aided with the identification of the correct *stx* subtype at the assembly level, due to the collection of *stx* sequence variants. However, nucleotide similarity amongst the sequences in stecodeDB would most likely cause issues for short-read mapping due to cross mapping. To test the impact of multiple "decoy" sequences, associated short reads of the STEC validation test set were mapped onto stecodeDB (with additional sequences of *recA* and *eae* from STEC O157 strain Sakai appended). Cross-mapping across intra-and-inter *stx* subtype sequences were observed (Figure SM3). When the mean read depth of all mapped *stx* sequences were normalised to the mean read depth of their respective *recA*, low normalisation were obtained with no values within their proximal, theoretical thresholds (i.e., proximal value of 1, 2 and 3 for single, double, and triple copies respectively; Figure SM3). Mapping to the entire stecodeDB would thus render inferences of multiple isogenic *stx* copies impossible.

To reduce the impact of these "decoy" sequences, consensus sequences were generated for *stx* subtypes within stecodeDB that had more than one sequence (henceforth referred to as stecodeCON). When the short reads were mapped to the stxrecaeae database (stecodeCON with *recA* and *eae* from STEC O157 strain Sakai appended), the normalisation values were relatively closer to their expected thresholds (Figure SM4A) but inter *stx* mapping was still observed Figure SM4). When coverage over the stxrecaeae database was interrogated, it was observed that mapping to the correct subtype will yield a coverage between 98.71% to 100% across the consensus sequence while mapping to the incorrect *stx* subtype will yield a coverage between 1.69 to 100% (Figure SM4B). The upper bound of 100% was due to P15-385, which had cross mapping with $stx_{2c}$. When this sample was omitted, this range changed to 1.69 to 97.34 %. For the two validation STEC genomes with a small truncation in *stx*, coverage was detected at 100% (Figure SM4B). To minimise the impact of cross mapping, all target sequences with a read coverage of greater than 98% were extracted, and mapping was performed on this reduced sequence set. This targeted mapping resulted in the correct re-call for majority of the samples (23/24, 95.83%), except for P15-385.

Results from targeted mapping also showed a general trend of increased ratios with majority of the double or triple copies being closer to their respective theoretical values (Figure SM4C). Exceptions were noted from the short-reads of RM10466, P15-385, and 95NR1 whereby the theoretical normalisation values were lower for the former two genomes and higher for the latter genome. In addition, FWSEC0006 and FWSEC0008 each had higher theoretical normalisation values for their single copy *stx* gene (Figure SM4C). This indicated that factors from either *in-vitro* growth, extraction or sequencing could impact on the normalisation as previously supposed (1), and it was prudent to include a buffer near the theoretical normalisation value of 2 to accommodate this. Based on the normalisation values, a threshold of greater than 2.1 was set as a compromise to infer a plausibility of multiple isogenic copies of the *stx* operon while a value between 2 and 2.1 would infer a possibility of multiple isogenic copies. (Figure SM4C).

#### General threshold for BLASTN-based inference of *stx* subtypes from short-read assemblies

Genomic similarities between *stx* subtypes could cause short-read assembly of *stx* genes to fail (i.e. non-contiguous assembly). To determine an appropriate threshold for BLASTN-based inferences of *stx* subtype using stecodeDB, the corresponding short reads from the validation set genomes were assembled (Table SM3) and *stx* subtypes were inferred on ABRicate using default settings. Of the 25 STEC genomes tested, eight had no *stx* subtypes detected while a discordant result was observed 95NR1 (Table SM4). The complete genome of 95NR1 had three copies of $stx_{2a}$, which in the process of short read assembly would have collapsed into a single copy.

The insertion of the IS*2* family insertion sequence within the *stx* operon (4) of STEC306, STEC307, and STEC308 resulted in non-contiguous assembly of their respective *stx* operons and thus, no *stx* could be detected with the default settings (Tables SM4 & SM5). The remaining five genomes that did not have *stx* genes detected under default settings were each localised to validation set genomes that each harboured either $stx_{2a}$, $stx_{2c}$ and $stx_{2d}$ subtypes in combination (Table SM4). The correct *stx* subtypes were detected in the short-read reassemblies of 19419 and RM10466, but the *stx* operon was not contiguously assembled and thus fell below the default detection threshold of ABRicate (Table SM4). Short-read reassembly of P15-385 only yielded a non-contiguously assembled $stx_{2a}$, while both FWSEC0008 and FWSEC0010 did not yield any significant *stx* assembly (Table SM5). This was despite having read coverage over the correct *stx* subtype genes in all three read sets (Figure SM4). It was highly likely that the high nucleotide identity between $stx_{2a}$ $stx_{2c}$ and $stx_{2d}$, in these three read sets, have impacted its assembly.

Based on the status of non-contiguous reassemblies of the test set (Table SM5), a BLASTN coverage threshold of greater than 21% (greater than 257 to 262 bases depending on *stx* subtype) was selected. This threshold allowed for the capture of the smallest assembled fragment in our validation dataset (last 276 bases for $stx_{2l}$) as well as being large enough to capture the smallest unit of the *stx* operon, the gene encoding the B subunit (size ranging from 263 bp to 269 bp based on sequences in stecodeDB). BLASTN inference (Table SM4) using this threshold, resulted in the correct recall of subtypes (excluding presence of multiple isogenic copies) of majority of the genomes tested (21/24, 87.5%). The only short-read reassembly that had detected multiple, isogenic *stx* copies with the new threshold was RM10466 (Table SM4).

### Tables

**Table SM1: Inter and intra BLASTN percentage identity amongst the 151 *stx* sequences in stecodeDB[a].**

| | Intra- *stx* subtype [a] | | Inter- *stx* subtype[b] | |
|---|---|---|---|---|
| subtype[a] | Minimum % ID | Maximum % ID | Minimum % ID | Maximum % ID |
| $stx_{1a}$ | 99.27 | 100 | 85.90 | 96.66 |
| $stx_{1c}$ | 99.59 | 100 | 86.65 | 96.66 |
| $stx_{1d}$ | 99.59 | 100 | 86.63 | 92.75 |
| $stx_{1e}$ | N/A | N/A | 85.90 | 86.97 |
| $stx_{2a}$ | 98.55 | 99.92 | 71.63 | 99.36 (95.81)[c] |
| $stx_{2b}$ | 97.34 | 99.92 | 70.96 | 94.81 |
| $stx_{2c}$ | 97.82 | 99.92 | 70.96 | 99.76 (95.25)[d] |
| $stx_{2d}$ | 96.87 | 99.92 | 71.05 | 99.76 (96.13)[e] |
| $stx_{2e}$ | 98.06 | 99.92 | 75.23 | 95.13 |
| $stx_{2f}$ | 99.19 | 99.84 | 70.96 | 76.15 |
| $stx_{2g}$ | 99.20 | 99.76 | 72.89 | 94.53 |
| $stx_{2h}$ | N/A | N/A | 71.88 | 93.36 |
| $stx_{2i}$ | N/A | N/A | 72.78 | 96.42 |
| $stx_{2j}$ | N/A | N/A | 72.61 | 91.01 |
| $stx_{2k}$ | 98.95 | N/A | 71.67 | 96.42 |
| $stx_{2l}$ | N/A | N/A | 72.78 | 95.81 |
| $stx_{2m}$ | 99.76 | 99.92 | 72.26 | 93.35 |
| $stx_{2n}$ | 99.92 | N/A | 73.45 | 91.42 |
| $stx_{2o}$ | N/A | N/A | 71.50 | 93.36 |

[a]: Self hits omitted from maximum % ID. If only one entry exists in database, both minimum and maximum % ID are listed as not applicable (N/A). If only two entries exist, only the maximum % ID is listed as N/A.

[b]: Inter-*stx* subtype comparisons were only performed within their *stx* type.

[c]: Listed in parenthesis is the maximum nucleotide identity with $stx_{2c}$ and $stx_{2d}$ omitted.

[d]: Listed in parenthesis is the maximum nucleotide identity with $stx_{2a}$ and $stx_{2d}$ omitted.

[e]: Listed in parenthesis is the maximum nucleotide identity with $stx_{2a}$ and $stx_{2c}$ omitted.

**Table SM2: Inference of stecodeDB sequences that do not have a representative subtype captured within VB01.**

| stecodeDB sequence | VB01 result | % coverage | % identity |
|---|---|---|---|
| KF926684_*stx*$_{1e}$ | stx$_{1c}$ | 99.92 | 86.97 |
| CP022279_*stx*$_{2h}$ | stx$_{2b}$ | 98.95 | 91.5 |
| FN252457_*stx*$_{2i}$ | stx$_{2e}$ | 98.62 | 94.24 |
| MZ571121_*stx*$_{2j}$ | stx$_{2d}$ | 99.11 | 89.94 |
| CP041435_*stx*$_{2k}$ | stx$_{2d}$ | 99.11 | 95.25 |
| KC339670_*stx*$_{2k}$ | stx$_{2d}$ | 100 | 96.13 |
| AM904726_*stx*$_{2l}$ | stx$_{2a}$ | 100 | 95.49 |
| ERR4297035_*stx*$_{2m}$ | stx$_{2a}$ | 99.27 | 92.86 |
| ERR4297034_*stx*$_{2m}$ | stx$_{2a}$ | 99.27 | 92.94 |
| OQ054797_*stx*$_{2m}$ | stx$_{2a}$ | 99.27 | 93.03 |
| CP113092_*stx*$_{2n}$ | stx$_{2d}$ | 99.19 | 88.17 |
| CP113094_*stx*$_{2n}$ | stx$_{2d}$ | 99.19 | 88.25 |
| MZ229604_*stx*$_{2o}$ | stx$_{2b}$ | 99.92 | 90.54 |

**Table SM3: Short-read reassembly of the validation test set.**

| Genome | Number of contigs | Cumulative assembly length (bases) | minimum contig length (bases) | maximum contig length (bases) | Average contig length (bases) | Assembly N50 (bases) |
|---|---|---|---|---|---|---|
| 194195 | 254 | 5439559 | 278 | 349083 | 21415.6 | 121029 |
| 95JB1 | 240 | 5244036 | 341 | 288493 | 21850.2 | 89036 |
| 95NR1 | 291 | 5277116 | 352 | 238414 | 18134.4 | 89036 |
| AUSMDU00002545 | 199 | 5392158 | 393 | 374711 | 27096.3 | 135186 |
| AUSMDU00014361 | 273 | 5466485 | 395 | 285510 | 20023.8 | 81718 |
| FWSEC0001 | 382 | 5319295 | 512 | 133719 | 13924.9 | 52328 |
| FWSEC0002 | 331 | 5242104 | 514 | 232742 | 15837.2 | 78938 |
| FWSEC0003 | 276 | 5204063 | 519 | 217663 | 18855.3 | 68610 |
| FWSEC0004 | 205 | 5218702 | 516 | 278761 | 25457.1 | 106497 |
| FWSEC0005 | 215 | 4893511 | 517 | 196664 | 22760.5 | 88973 |
| FWSEC0006 | 229 | 5113063 | 515 | 308133 | 22327.8 | 96552 |
| FWSEC0007 | 247 | 5195925 | 515 | 280186 | 21036.1 | 91546 |
| FWSEC0008 | 170 | 4892363 | 528 | 278949 | 28778.6 | 73235 |
| FWSEC0009 | 266 | 5230005 | 518 | 196934 | 19661.7 | 53676 |
| FWSEC0010 | 131 | 4985338 | 528 | 289184 | 38056 | 167402 |
| MBT-5 | 279 | 5506343 | 346 | 252268 | 19736 | 97406 |
| P15-385 | 152 | 5048765 | 335 | 295266 | 33215.6 | 125198 |
| PNUSAE009425 | 60 | 5078807 | 482 | 420821 | 84646.8 | 165550 |
| RM10466 | 100 | 5189638 | 517 | 495832 | 51896.4 | 186672 |
| STEC2017-197 | 122 | 5013514 | 316 | 699117 | 41094.4 | 147678 |
| STEC306 | 110 | 5104087 | 379 | 254307 | 46400.8 | 145385 |
| STEC307 | 120 | 5104356 | 337 | 285780 | 42536.3 | 145385 |

| Genome | Number of contigs | Cumulative assembly length (bases) | minimum contig length (bases) | maximum contig length (bases) | Average contig length (bases) | Assembly N50 (bases) |
|---|---|---|---|---|---|---|
| STEC308 | 120 | 5104055 | 342 | 254307 | 42533.8 | 140716 |
| STEC865 | 74 | 4857935 | 375 | 371697 | 65647.8 | 195020 |
| STEC866 | 75 | 4858112 | 366 | 371697 | 64774.8 | 195020 |

**Table SM4: Inferences of *stx* subtype from short-read reassemblies of the STEC validation dataset.**

| | | ABRicate inferred *stx* subtypes | |
|---|---|---|---|
| **Genome** | ***stx* subtype[a]** | **Default setting** | **Revised setting** |
| 194195 | $stx_{2a}$, $stx_{2c}$ | Not detected | $stx_{2a}$, $stx_{2c}$ |
| 95JB1 | $stx_{1a}$, $stx_{2a}$ | $stx_{1a}$, $stx_{2a}$ | $stx_{1a}$, $stx_{2a}$ |
| 95NR1 | $stx_{1a}$, $stx_{2a}$, $stx_{2a}$, $stx_{2a}$ | $stx_{1a}$, $stx_{2a}$ | $stx_{1a}$, $stx_{2a}$ |
| AUSMDU00002545 | $stx_{1a}$, $stx_{2c}$ | $stx_{1a}$, $stx_{2c}$ | $stx_{1a}$, $stx_{2c}$ |
| AUSMDU00014361 | $stx_{2a}$ | $stx_{2a}$ | $stx_{2a}$ |
| FWSEC0001 | $stx_{1a}$, $stx_{2a}$ | $stx_{1a}$, $stx_{2a}$ | $stx_{1a}$, $stx_{2a}$ |
| FWSEC0002 | $stx_{1a}$ | $stx_{1a}$ | $stx_{1a}$ |
| FWSEC0003 | $stx_{1a}$ | $stx_{1a}$ | $stx_{1a}$ |
| FWSEC0004 | $stx_{1a}$, $stx_{2a}$ | $stx_{1a}$, $stx_{2a}$ | $stx_{1a}$, $stx_{2a}$ |
| FWSEC0005 | $stx_{1a}$, $stx_{2a}$ | $stx_{1a}$, $stx_{2a}$ | $stx_{1a}$, $stx_{2a}$ |
| FWSEC0006 | $stx_{2a}$ | $stx_{2a}$ | $stx_{2a}$ |
| FWSEC0007 | $stx_{1a}$ | $stx_{1a}$ | $stx_{1a}$ |
| FWSEC0008 | $stx_{2a}$, $stx_{2d}$, $stx_{2d}$ | Not detected | Not detected |
| FWSEC0009 | $stx_{2a}$ | $stx_{2a}$ | $stx_{2a}$ |
| FWSEC0010 | $stx_{2a}$, $stx_{2d}$ | Not detected | Not detected |
| MBT-5 | $stx_{1a}$, $stx_{2a}$ | $stx_{1a}$, $stx_{2a}$ | $stx_{1a}$, $stx_{2a}$ |
| P15-385 | $stx_{2a}$, $stx_{2d}$, $stx_{2d}$ | Not detected | $stx_{2a}$ |
| PNUSAE009425 | $stx_{2n}$ | $stx_{2n}$ | $stx_{2n}$ |
| RM10466 | $stx_{2a}$, $stx_{2a}$, $stx_{2d}$ | Not detected | $stx_{2a}$, $stx_{2a}$, $stx_{2d}$ |
| STEC2017-197 | $stx_{2d}$, $stx_{2d}$ | $stx_{2d}$ | $stx_{2d}$ |
| STEC306 | $stx_{2l}$ (disrupted) | Not detected | $stx_{2l}$ |
| STEC307 | $stx_{2l}$ (disrupted) | Not detected | $stx_{2l}$ |
| STEC308 | $stx_{2l}$ (disrupted) | Not detected | $stx_{2l}$ |
| STEC865 | $stx_{1c}$ | $stx_{1c}$ | $stx_{1c}$ |
| STEC866 | $stx_{1c}$ | $stx_{1c}$ | $stx_{1c}$ |
| MG1655 | Not STEC | Not detected | Not detected |
| EC958 | Not STEC | Not detected | Not detected |
| E2348/69 | Not STEC | Not detected | Not detected |

**Table SM5: Non-contiguous assembly of the *stx* operon following short-read reassembly in eight STEC validation test set.**

| Genome | stx content | Contig(s) where stx was assembled in | | Best stecodeDB hit | | Coverage | | % coverage | |
|---|---|---|---|---|---|---|---|---|---|
| | | First contig | Second contig | First contig | Second contig | First contig | Second contig | First contig | Second contig |
| 194195 | 2a | Contig_104 | Contig_148 | AF461171_stx2a | AB030484_stx2a | 1-731/1241 | 754-1241/1241 | 58.9 | 39.32 |
| | 2c | Contig_73 | - | AM982821_stx2c | - | 754-1241/1241 | - | 39.32 | - |
| FWSEC0008 | 2a | Contig_153 | Contig_154 | FM998851_stx2a | FM998851_stx2a | 1-92/1241 | 1121-1241/1241 | 7.41 | 9.75 |
| | 2d | Not assembled | Not assembled | Not applicable | Not applicable | Not applicable | Not applicable | Not applicable | Not applicable |
| | 2d | Not assembled | Not assembled | Not applicable | Not applicable | Not applicable | Not applicable | Not applicable | Not applicable |
| FWSEC0010 | 2a | Not assembled | Not assembled | Not applicable | Not applicable | Not applicable | Not applicable | Not applicable | Not applicable |
| | 2d | Not assembled | Not assembled | Not applicable | Not applicable | Not applicable | Not applicable | Not applicable | Not applicable |
| P15-385 | 2a | Contig_59 | Contig_83 | GQ429170_stx2a | FM998851_stx2a | 790-1241/1241 | 1-731/1241 | 36.42 | 58.9 |
| | 2d | Not assembled | Not assembled | Not applicable | Not applicable | Not applicable | Not applicable | Not applicable | Not applicable |
| | 2d | Not assembled | Not assembled | Not applicable | Not applicable | Not applicable | Not applicable | Not applicable | Not applicable |
| RM10466 | 2a | Contig_54 | - | GQ429163_stx2a | - | 738-1241/1241 | - | 40.61 | - |
| | 2a | Contig_55 | - | GQ429170_stx2a | - | 738-1241/1241 | - | 40.61 | - |
| | 2d | Contig_11 | - | AF500190_stx2d | - | 609-1241/1241 | - | 51.01 | - |
| STEC306 | 2l | Contig_55 | Contig_80 | AM904726_stx2l | AM904726_stx2l | 1-963/1241 | 965-1241/1241 | 77.6 | 22.32 |
| STEC307 | 2l | Contig_44 | Contig_86 | AM904726_stx2l | AM904726_stx2l | 1-963/1241 | 965-1241/1241 | 77.6 | 22.32 |
| STEC308 | 2l | Contig_59 | Contig_86 | AM904726_stx2l | AM904726_stx2l | 1-963/1241 | 965-1241/1241 | 77.6 | 22.32 |

### Figures

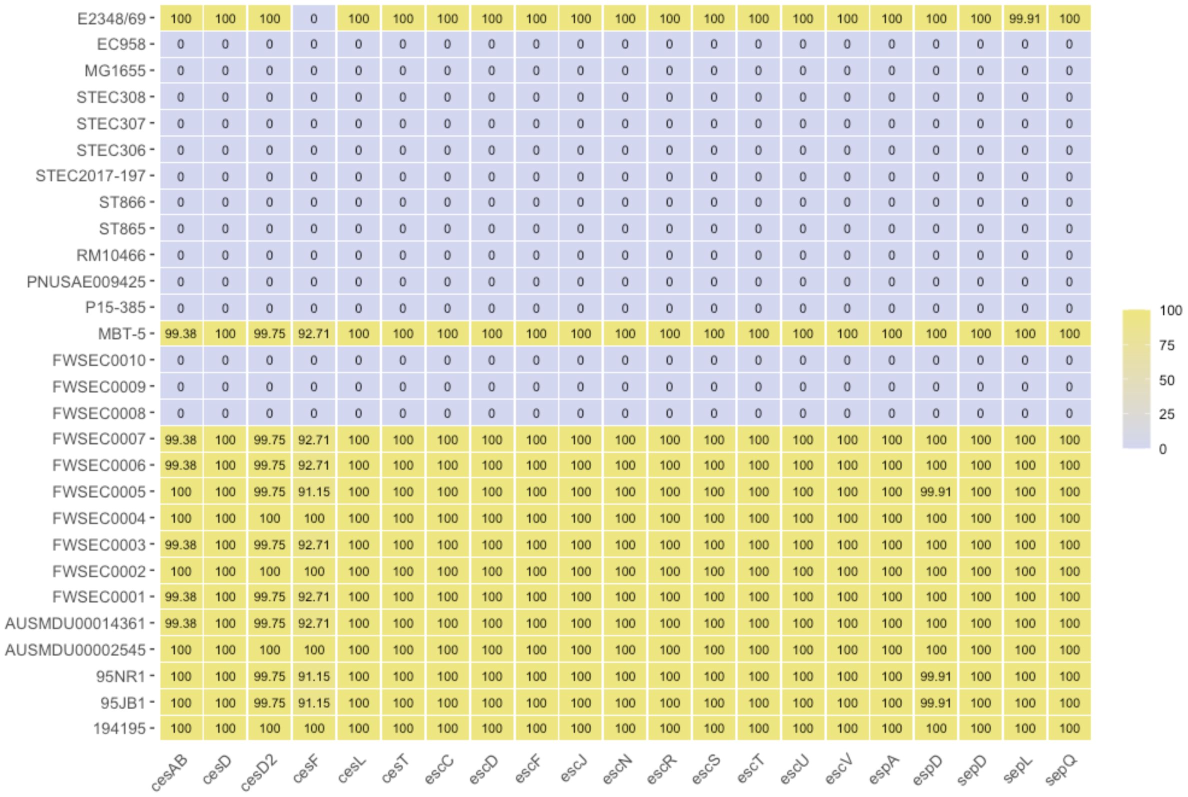


**Figure SM1: Presence of key LEE genes within the validation test set represented as a heatmap.** Genes listed here encode for the chaperones, secretion machinery (both Esc and Sep), and translocator proteins of the LEE. Queries were made using ABRicate against the VFDB database packaged within. Numbers within each tile represent proportion (in percentage) of the gene covered and colour coded according to the gradient bar. Image was generated using ggplot2 version 3.4.2. (5)

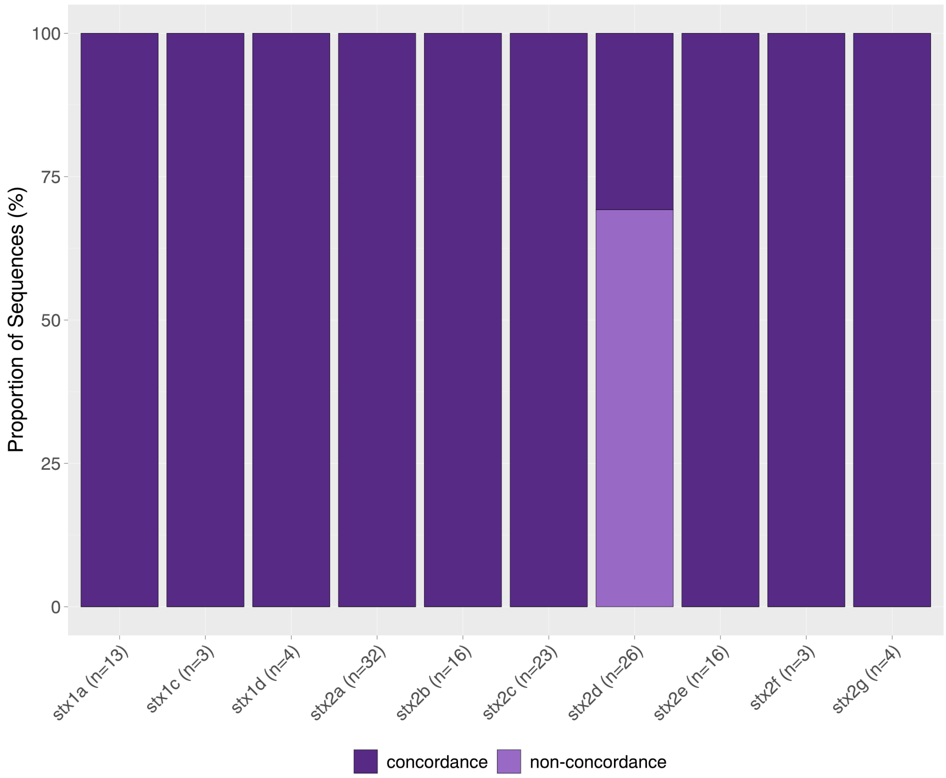


**Figure SM2: Proportion of concordance when stecodeDB sequences were checked against the VB01 database.** Concordance status of the barplot is colour coded according to the figure key. Barplot was generated using ggplot version 3.4.2. (5)

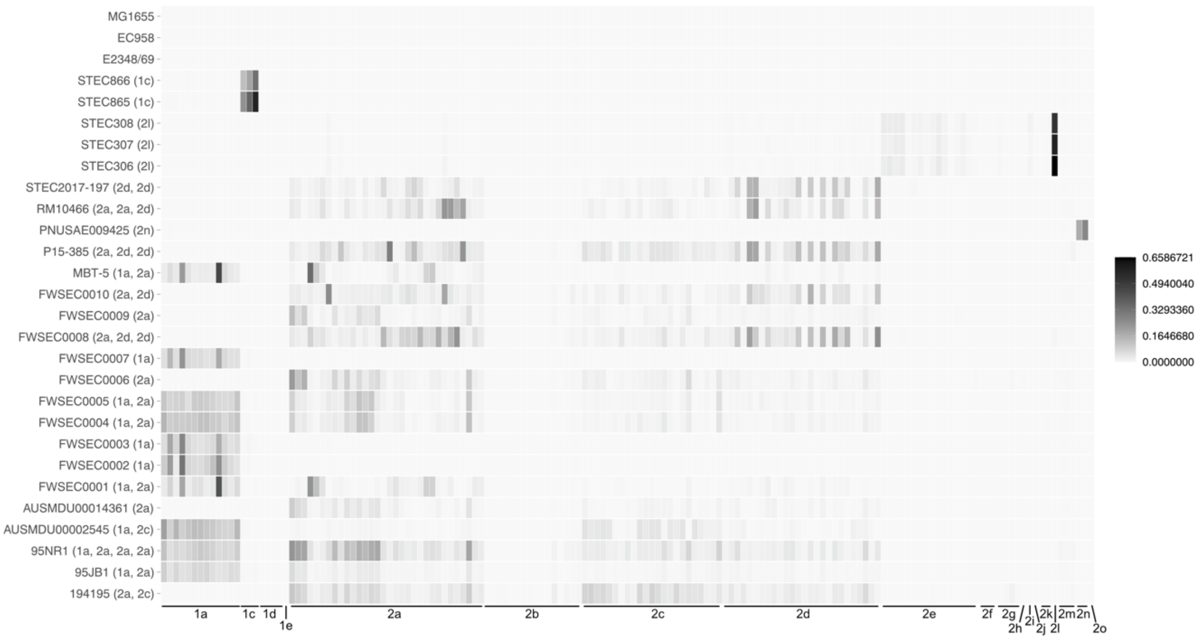


**Figure SM3: Heatmap for inference of multiple copies of isogenic *stx* using the entire stecodeDB.** Each sequence in stecodeDB is represented as a single rectangle and grouped accordingly to their *stx* subtype. Normalisation values of each *stx* sequences was scaled according to the gradient bar. Heatmap was generated using ggplot2 version 3.4.2. (5).

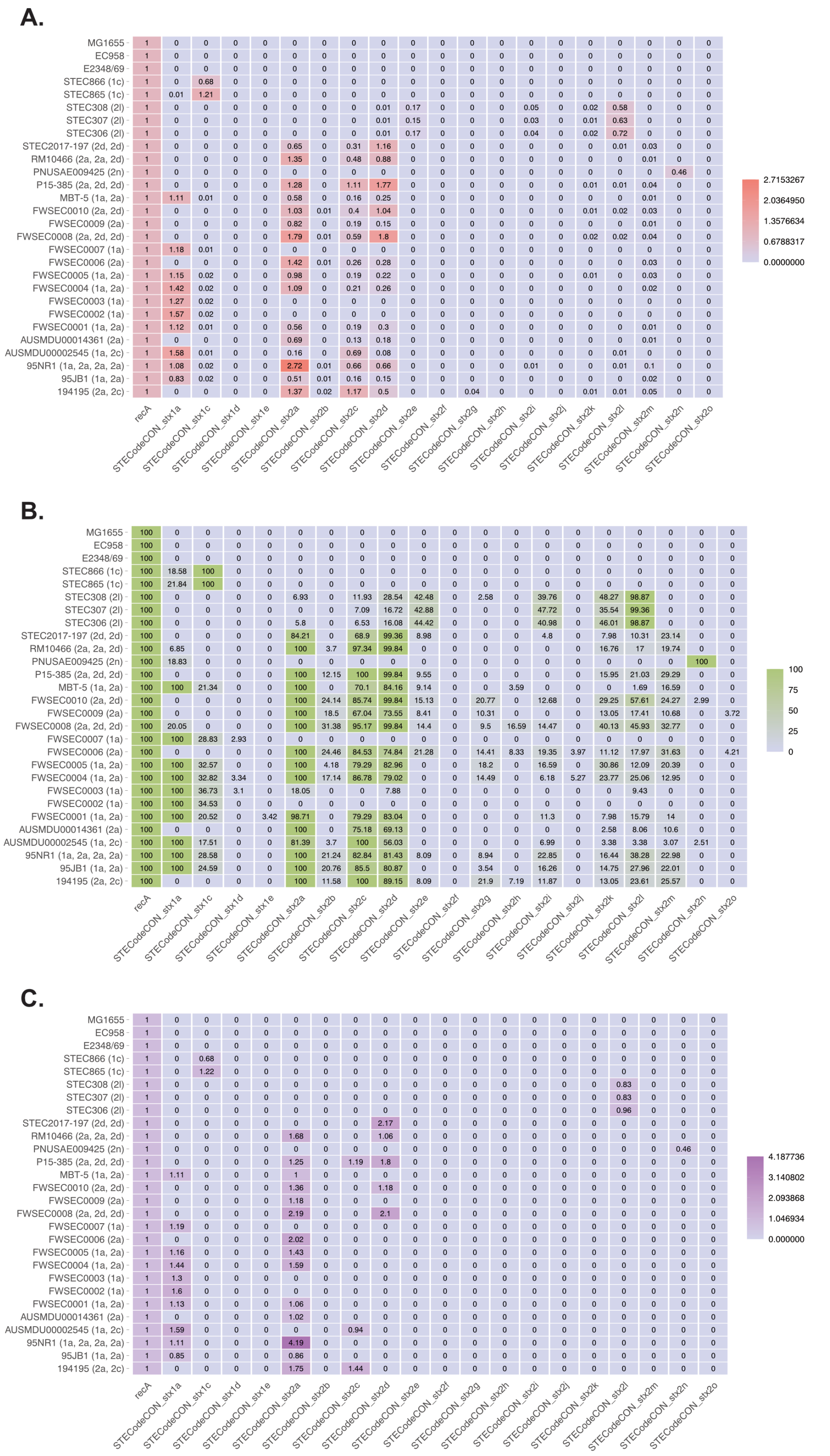


**Figure SM4: Heatmaps for inferences of multiple isogenic *stx* copies.** (A) Normalisation values of *stx* sequences when short reads of the STEC validation set were mapped onto the sequences in the stxrecaeae database. (B) Percentage coverage of sequences when short reads were mapped onto stxrecaeae. (C) Normalisation values of stx sequences after targeted mapping. Values in all panels are colour coded according to their respective gradient bar. Heatmaps were generated using ggplot2 version 3.4.2. (5)
